# NanoBRET Tracer Development for Class I Bromodomain Target Engagement in Live Cells

**DOI:** 10.1101/2025.11.25.689777

**Authors:** Molly S. Sneddon, Chun-Ju Tsou, Xiang Fu, Anang A. Shelat, William C. K. Pomerantz

## Abstract

Epigenetic reader proteins, such as bromo-domains, are often associated with diseases such as cancer and inflammation. BET bromodomain inhibitors have been studied extensively; however, non-BET bromodomains are understudied. Moreover, available high-throughput biological assays to assess inhibitors are limited. One non-BET bromodomain-containing protein, BPTF, has a recently reported inhibitor, BZ1, with an in vitro affinity of 6.3 nM. Additionally, BZ1 is known to be non-selective towards other class I bromodomains PCAF, GCN5, and CECR2. Here, we use a BZ1 analog, BZ1-THQ, to design a small molecule NanoBRET tracer, **MS-1**, for assessing inhibitor functional activity through live-cell target engagement against the BPTF bromodomain. Further, we investigate the versatility of **MS-1** against PCAF, GCN5, and CECR2. We observe that **MS-1** is a broadly applicable NanoBRET tracer for class I bromodomains, effectively binding BPTF, PCAF, GCN5, and CECR2 in HEK293T cells at low to sub-micromolar concentrations. We report EC_50_ values of commercially available and in-house inhibitors to demonstrate tracer versatility for future target engagement studies and inhibitor development.

**Table of Contents artwork:** 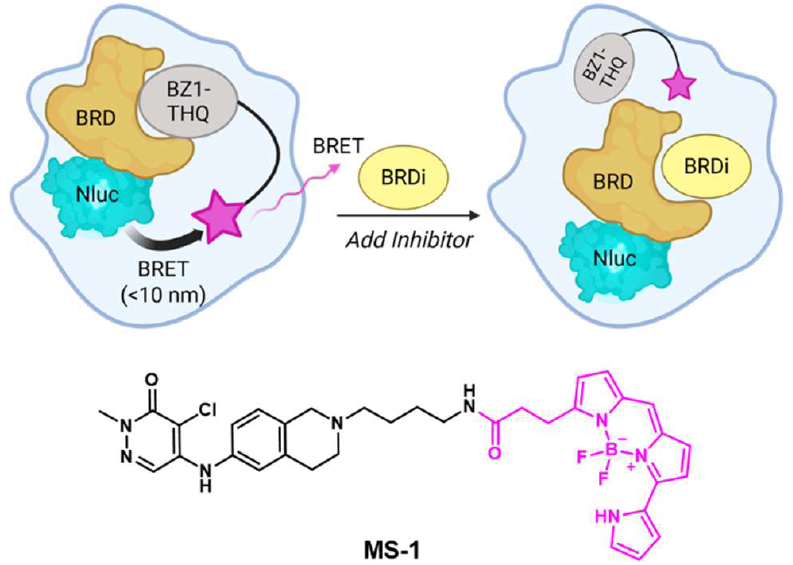

## INTRODUCTION

Bromodomains belong to a class of epigenetic “reader” domains which recognize histone post-translational modifications on chromatin and dictate downstream genetic processes.^1^ There are 61 human bromodomains found in 46 distinct proteins split among eight classes. Bromodomains recognize acetylated lysine residues most notably on histone proteins. While bromodomains are important for regular function of cells such as myogenesis, neuroplasticity, and cell-cycle regulation, dysregulation of bromodomains can result in a myriad of diseases including cancer and inflammation.^2–6^ Given their role in disease, development of inhibitors to prevent bromodomains from interacting with chromatin and subsequently decreasing aberrant gene expression is an active area of study.

Cellular target engagement is an important consideration for drug development for bridging the gap between in vitro and in vivo studies of bromodomain-containing proteins. While commonly used PAMPA (parallel artificial membrane permeability assay) and Caco-2 assays can assess compound permeability and efflux liabilities, target engagement assays provide a combined readout of cellular permeability and functional effects. One assay which can be used to determine bromodomain inhibitor engagement in live cells is known as NanoBRET based on bioluminescent resonance energy transfer. NanoBRET requires a nanoluciferase (Nluc) enzyme to be fused to a protein of interest at either the N- or C-terminus of the protein and expressed in cells. Secondly, a small molecule tracer must be developed which contains the protein ligand, linker, and fluorophore. BODIPY 576/589 is a commonly used fluorophore for NanoBRET assays. The ligand binds to the protein, and the BODIPY fluorophore has a structural requirement to be in proximity of less than 10 nm to the Nluc enzyme. Finally, when the Nluc substrate furimazine is added, Nluc oxidatively converts the substrate into furimamide, CO_2_, and light.^7,8^ The energy from the bioluminescence can be transferred to the BODIPY fluorophore leading to the BRET signal. When unlabeled bromodomain inhibitors are added, the tracer is displaced resulting in a dose-dependent reduction in BRET (**Figure 1A**).

**Figure 1.**
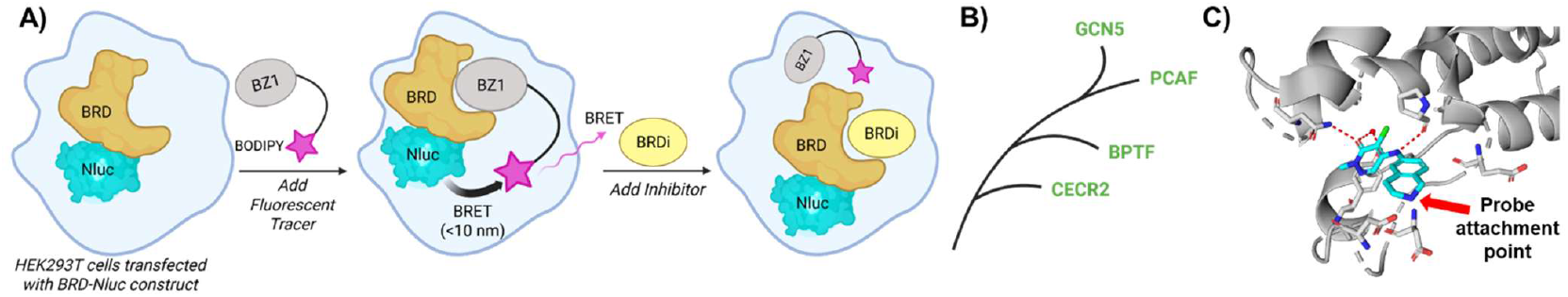
**A)** Graphical representation of the NanoBRET platform where a bromodomain inhibitor displaces a chemical tracer molecule in HEK293T cells and allows for the assessment of cellular permeability via target engagement. **B)** The class I bromodomain structural homology tree branch including BPTF, PCAF, CECR2, and GCN5. **C)** Co-crystal structure of BZ1-THQ (blue) with the BPTF-BRD (gray) (PDB 7RWQ).

Class II BET (bromodomain and extra-terminal) bromo-domains are well-characterized and have commercial tracers available to study inhibitor engagement in live cells. Novel tracers have also been developed for screening such as JQ1-NCT described by Bradner and co-workers.^9^ We have become interested in class I bromodomains, particularly for developing bromodomain inhibitors of BPTF (bromodomain and PHD finger transcription factor). BPTF is an emerging cancer target in part due to regulation of Myc signaling genes. Additional Class I bromodomains, PCAF, CECR2, and GCN5, have also been targeted for diseases such as HIV and glioblastoma, breast cancer, and osteoporosis, respectively (**Figure 1B**).^10–13^ Inhibitor development against class I bromodomains is limited by fewer standardized assays to study compound engagement in cells. Cellular engagement thermal shift assay (CETSA) is one such assay, however some proteins like BPTF are very large (>250 KDa), and western blotting for stabilized protein is both low throughput and challenging.^14^ Additionally, heating cells can cause changes to binding equilibria and cellular permeability, and weaker interactions may not result in measurable stabilization.^15^ A promiscuous non-BET bromodomain inhibitor known as bromosporine was previously developed by Picaud et al. and might initially be considered for an in-cell bromodomain tracer.^16^ Promiscuous inhibitors are attractive as they can be used to develop target engagement assays against multiple proteins. However, we have shown bromosporine to bind to BPTF weakly in vitro (K_d_ = 37 μM) and would thus be a poor candidate for tracer development.^17^

While a NanoBRET assay has been previously reported for BPTF, constructs and optimized conditions are not well described nor have their broader utility towards other class I bromodomains been evaluated.^18^ Previously, Zahid et al. developed a BPTF bromodomain (BPTF-BRD) inhibitor known as BZ1 with an affinity of 6.3 nM. BZ1 was shown to preferentially bind to the BPTF-BRD and demonstrated additional off-target activity for the other class I bromodomains with affinities ranging from 11-44 nM.^19^ BZ1 also inhibits class IV bromodomains BRD7 and BRD9. Here, we use a BZ1 analog as a promiscuous class I bromodomain inhibitor to develop a series of NanoBRET tracers targeting class I bromodomains. Our optimal tracer was first characterized against the BPTF-Nluc bromodomain constructs using a suite of established inhibitors from the literature. Subsequently, we compare our results with Caco-2 permeability data for a series of BPTF bromodomain inhibitors, including several new analogs. Finally, we characterized our tracer against additional bromodomains CECR2, PCAF, and GCN5. These studies yield a well-characterized NanoBRET tracer that can be used to assess inhibitor engagement and permeability for all class I bromodomains. Such a tracer may also be useful for additional bromodomain-containing proteins, such as BRD7 and BRD9, which are important anticancer targets.^20,21^

## RESULTS AND DISCUSSION

To design a versatile class I bromodomain NanoBRET tracer, a BZ1 tetrahydroisoquinoline derivative (BZ1-THQ) was chosen due to its increased cellular permeability over BZ1. Additionally, the BZ1-THQ amine exit vector has been previously studied through analysis of structure-activity relationship data and protein co-crystal structures (**Figure 1C**).^22^ Information regarding the exit vector for linker and BODIPY dye attachment is necessary for development of effective NanoBRET tracers. Three NanoBRET tracers (**MS-1, MS-2**, and **MS-3**) were synthesized with the BZ1-THQ derivative as the bromodomain binding compound, BODIPY 576/589 as the fluorophore, and three different linkers of varying length and composition (**Figure 2**).

**Figure 2.**
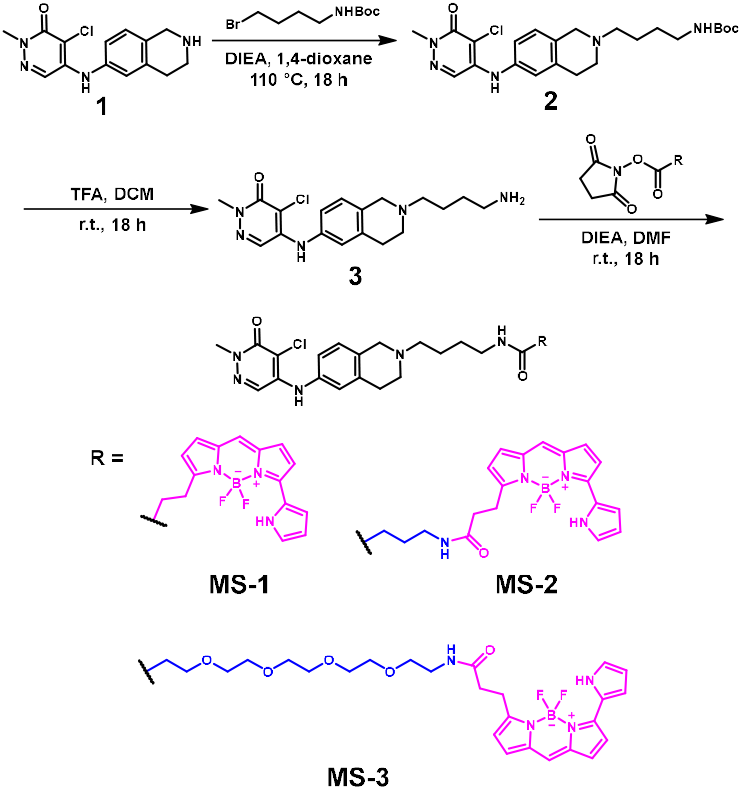
Synthetic scheme and structures of the three tracers, **MS-1, MS-2**, and **MS-3**.

The tracers were first assessed for their ability to engage the BPTF-BRD-Nluc constructs in HEK293T cells and induce a large BRET response. Given NanoBRET is a proximity-based assay, the position of the BODIPY fluorophore relative to the Nluc enzyme must be optimized. The Nluc enzyme was placed at either the N- or C-terminus of the BPTF-BRD (indicated as Nluc-BPTF-BRD and BPTF-BRD-Nluc, respectively). **MS-1, MS-2**, and **MS-3** were then titrated against both BPTF-BRD constructs to determine which provided the widest assay window as the optimal tracer-construct combination (**Figure 3A, S1**). The titration experiments indicated that all three tracers demonstrated some level of engagement with each BPTF-BRD construct. **MS-1**, which had no additional linker, possessed the highest affinity (0.6 ± 0.1 μM) and resulted in the widest assay window (60 milliBRET units, mBu) for the BPTF-BRD-Nluc construct. Tracer engagement decreased as linker length increased with **MS-2** and **MS-3** having lower affinities and assay windows. The Nluc-BPTF-BRD construct resulted in similar engagement with each tracer with slightly smaller assay windows. A similar assay window between constructs could be rationalized based on the small size of the BPTF-BRD (14 KDa). Additionally, the N- and C-termini of bromodomains are often in close proximity and could engage with the BODIPY fluorophore similarly. Thus, **MS-1** and the BPTF-BRD-Nluc construct were chosen as the optimal combination for subsequent competition experiments with bromodomain inhibitors.

**Figure 3.**
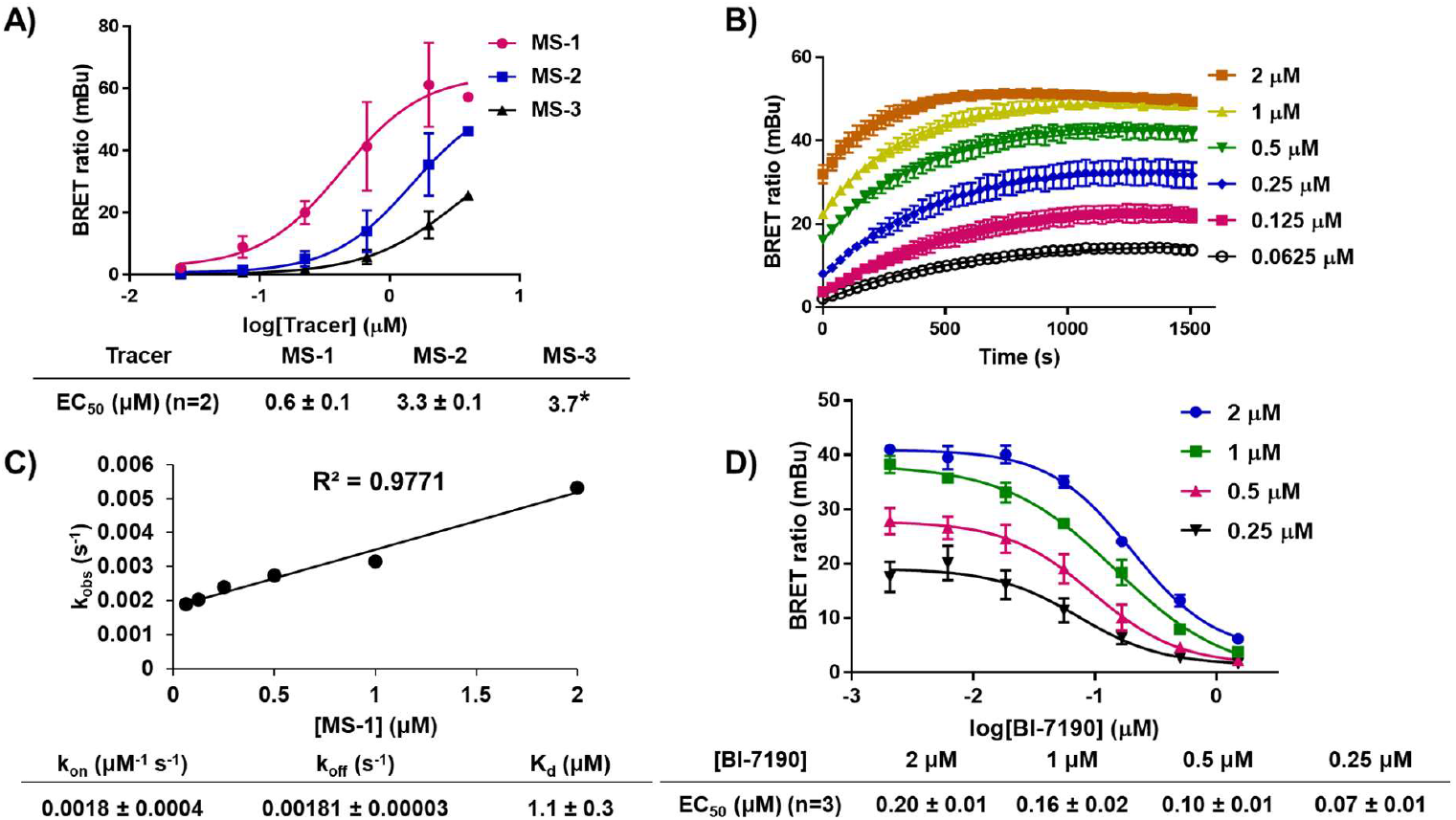
**A)** Tracer titration plots against BPTF-BRD-Nluc in HEK293T cells and their EC_50_values. *second replicate EC_50_ could not be determined due to an incomplete curve. **B)** Kinetics data with **MS-1** at varying concentrations over 25 minutes (truncated from 1 hour). **C)** Linear plot of k_obs_ values at each concentration of **MS-1**. The k_on_ and k_off_ values were determined from the linear fit equation and were used to calculate the K_d_. **D)** BI-7190 titration plots at various **MS-1** concentrations to determine ideal assay window. EC_50_ values of BI-7190 are reported at various **MS-1** concentrations. All graphical data represents mean ± standard deviation where **A, B**, and **C** are technical duplicates with n=2, and **D** is reported as a technical duplicate with n=3.

Before testing inhibitors, the K_d_ value of the tracer with the BPTF-BRD was measured kinetically by determining the k_on_ and k_off_ rates of the NanoBRET tracer. The K_d_ of the tracer enables the conversion of EC_50_ values to apparent K_i_ values. Varying concentrations of **MS-1** were dosed, and measurements were taken every 35 seconds over the course of one hour (**Figure 3B**). The association of **MS-1** with the BPTF-BRD was fast (~ 15 minutes). Data was fit using one-phase association and k_obs_ values were determined for each concentration. The k_obs_ values were then fit to a linear curve (**Figure 3C**) which was used to derive the k_on_ and k_off_ values resulting in a K_d_ of 1.1 ± 0.3 μM. This affinity is within two-fold of the EC_50_ derived for **MS-1**, indicating good cell permeability of the tracer. Additionally, this moderate affinity is an ideal range for future competition experiments.

One additional assessment was conducted where a 25X molar excess of the N-methylated version of BZ1-THQ, compound **4** (**Figure 5B**), was added to a titration of tracer to ensure **MS-1** could be competed away. This experiment ensures the tracer engages in a low level of non-specific interactions with the BPTF-BRD. **MS-1** could be largely competed away by **4**, indicating the tracer exhibits low levels of non-specific interactions (**Figure S2**). Overall, **MS-1** was demonstrated to be an effective NanoBRET tracer for the BPTF-BRD.

**MS-1** was next assessed for its utility in a competition-based NanoBRET experiment for the determination of live cell bromodomain engagement. The first compound tested, BI-7190, is a commercially available BPTF-BRD inhibitor that is both cell permeable and possesses a high affinity (ITC K_d_ = 85 nM).^23^ BI-7190 was titrated against varying concentrations of **MS-1** to determine the minimum tracer concentration required for an assay window above two-fold mBu and within two-fold of the BI-7190 affinity (**Figure 3D**).^24^ Each concentration of **MS-1** was able to be competed away by BI-7190. A concentration of 0.5 μM **MS-1** gave an ideal assay window approaching 30 mBu and was below the tracer EC_50_ as recommended by the Promega technical manual.^24,25^ The scope of **MS-1** efficacy across a variety of commercially available inhibitors was then examined as shown in **Figure 4**. NVS-BPTF-1, TP-238, and DC-BPi-11 all showed engagement with the BPTF-BRD. DC-BPi-11 and BI-7190 closely corroborated reported literature values for affinity with the BPTF-BRD.^23,26,27^ NVS-BPTF-1 and TP-238 were notably weaker than reported in vitro values which could be attributed to the poor solubility of NVS-BPTF-1 and poor cellular permeability of TP-238 based on reported PAMPA data (**Figure 4B**). S-AU1 and AU1 (Rac) were also tested, however neither showed full inhibition of the BPTF-BRD which could also be attributed to the known poor solubility and weaker affinity of AU1 (K_d_ = 2.8 μM, **Figure S3**).^28^

**Figure 4.**
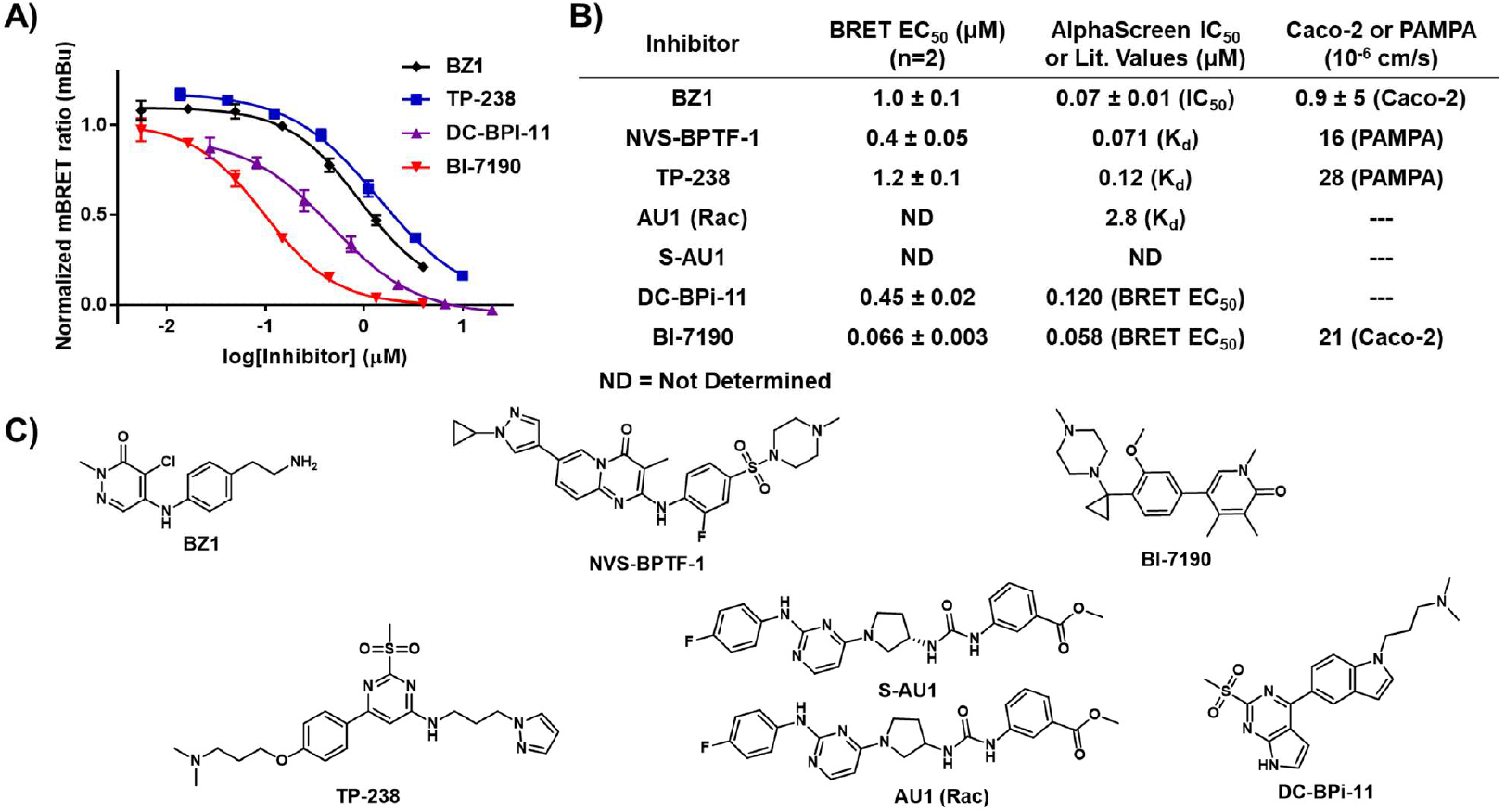
**A)** Titration plots of commercial and in-house inhibitors of the BPTF-BRD-Nluc with 0.5 μM **MS-1. B)** Experimentally determined BRET EC_50_ values and literature affinity and permeability data for each inhibitor against the BPTF-BRD. **C)** Structures of each tested inhibitor. NanoBRET EC_50_ values are mean ± standard deviation reported as technical triplicates with n=2.

As a final test case for the BPTF-BRD, we evaluated three novel BPTF-BRD inhibitors based on BZ1. **MS-1** was used to assess engagement of **4, 5**, and **6** against the BPTF-BRD (**Figure 5**). Each compound engaged the BPTF-BRD, and the affinities corroborated both the in vitro AlphaScreen competitive inhibition data as well as Caco-2 permeability data. Compound **4** exhibited the highest in vitro affinity as well as the highest in cellulo affinity with an EC_50_ of 0.9 ± 0.2 μM. Compound **5** demonstrated a slightly lower EC_50_ of 1.4 ± 0.2 μM which corroborated its lower in vitro affinity. **5** exhibits a higher permeability, and this validates the 1.5-fold lower affinity in cells compared to the 2.5-fold lower affinity in vitro. Compound **6** demonstrated the lowest affinity at 4.0 ± 0.3 μM in cells which aligned with the in vitro affinity of 2.1 μM.

**Figure 5.**
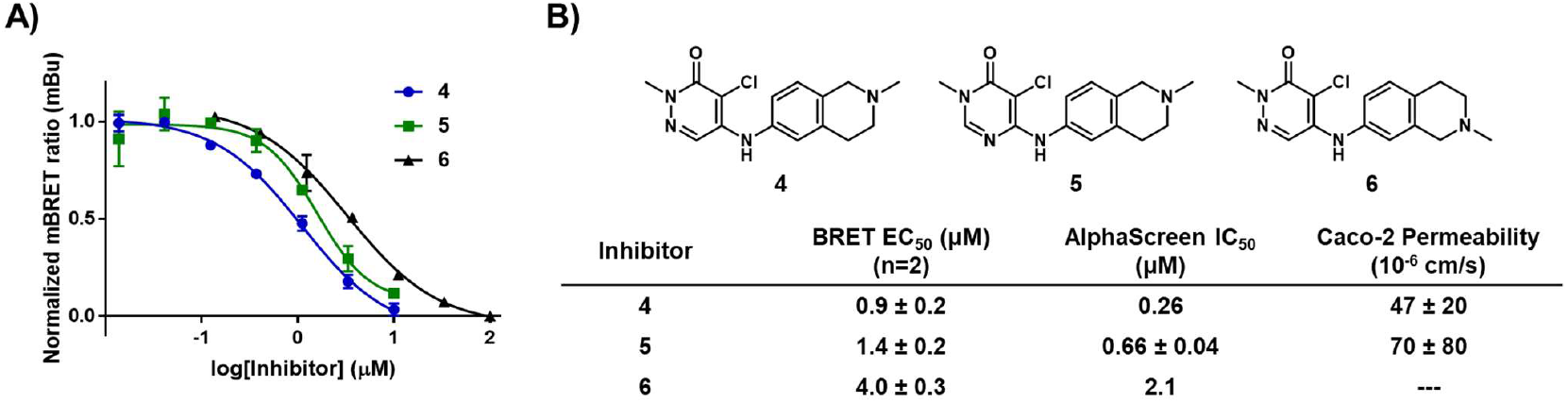
**A)** Titration plots of **4, 5, 6** against BPTF-BRD-Nluc using 0.5 μM **MS-1. B)** Structures of tested compounds and determined BRET EC_50_ values. In vitro AlphaScreen IC_50_ values and Caco-2 permeability are included for comparison for compounds **4** and **5**. Caco-2 permeability was not determined for compound **6**. NanoBRET EC_50_ values are mean ± standard deviation reported as technical triplicates with n=2.

Given the encouraging results of **MS-1** as a NanoBRET tracer for the BPTF-BRD, we next evaluated its versatility as a tracer against additional class I bromodomains. First, engagement of **MS-1** with full-length PCAF was investigated. **MS-1** was titrated against two constructs of full-length PCAF containing either N- or C-terminal Nluc to determine the optimal position of the Nluc enzyme. The C-terminal PCAF-Nluc construct showed significantly better engagement with **MS-1** compared to Nluc-PCAF (**Figure S4**). When titrated against 4, **MS-1** was competed away, indicating a low level of non-specific interactions with PCAF-Nluc (**Figure S4**). A concentration of 1 μM **MS-1** was chosen for an assay window approaching 20 mBu and was below the tracer EC_50_ against PCAF-Nluc. Subsequently, the commercially available PCAF inhibitor L-Moses was titrated against PCAF-Nluc and resulted in an EC_50_ of 0.28 ± 0.05 μM which is comparable to the literature K_d_ value of 0.126 μM.^29^ BZ1 was also tested and exhibited an EC_50_ of 3.0 ± 0.4 μM (**Figure 6A**). Given the tracer engages PCAF readily in cells and can validate previous data obtained for literature and in-house compounds, we conclude that **MS-1** can also be used for rank ordering PCAF inhibitor affinities in live cells.

**Figure 6.**
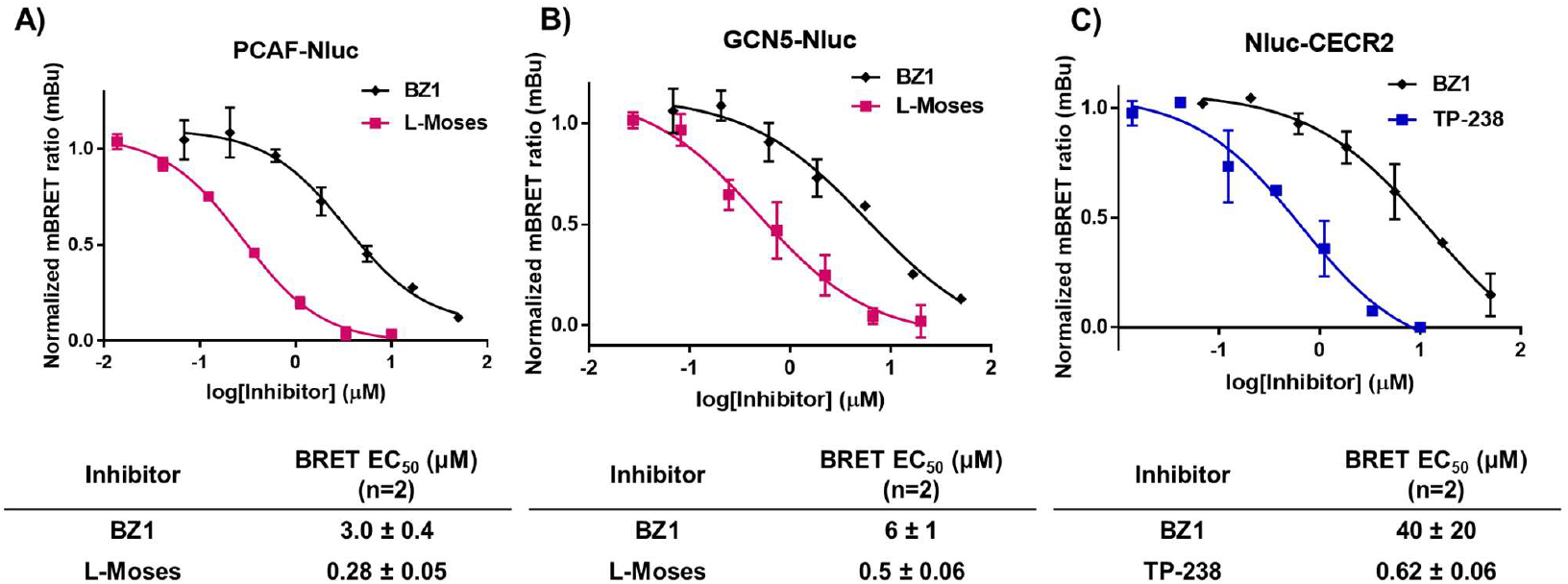
Titration plots and experimentally determined BRET EC_50_ values for three different classI bromodomain-containing proteins. **A)** BZ1 and L-Moses against PCAF-Nluc using 1 μM **MS-1**.B) BZ1 and L-Moses against GCN5-Nluc using 1 μM **MS-1. C)** BZ1 and TP-238 against Nluc-CECR2 using 0.5 μM **MS-1**. NanoBRET EC_50_ values are mean ± standard deviation reported as technical triplicates with n=2.

GCN5 was next examined for its ability to be engaged by **MS-1**. The C-terminal full-length GCN5-Nluc construct was titrated against **MS-1** and exhibited dose response. A concentration of 1 μM **MS-1** gave an assay window approaching 20 mBu and was below the tracer EC_50_ value against GCN5-Nluc. The non-specificity assessment with **4** indicated that **MS-1** could also be competed away similar to our results with PCAF supporting low levels of non-specific interactions with GCN5-Nluc (**Figure S5**). The commercial inhibitor L-Moses was titrated first against GCN5-Nluc and resulted in an EC_50_ of 0.50 ± 0.06 μM. BZ1 was then titrated against GCN5-Nluc, and it exhibited a weak affinity of 6 ± 1 μM (**Figure 6B**). This weaker affinity was expected according to the previously determined BZ1 in vitro affinity.

Lastly, we evaluated **MS-1** against the final class I bromo-domain CECR2. **MS-1** was titrated against the N-terminal full length Nluc-CECR2. Similar to our prior results, **MS-1** also engaged Nluc-CECR2. A concentration of 1 μM **MS-1** was chosen for an assay window of around 20 mBu and was below the tracer EC_50_ value against Nluc-CECR2. The non-specificity assessment against **MS-1** with **4** showed less-efficient competition compared to the other bromodomains, but this was likely due to the inherent lower affinity of **4** with CECR2 (**Figure S6**). The commercial inhibitor TP-238 was first titrated against CECR2 with an **MS-1** concentration of 1 μM, and an EC_50_ of 0.62 ± 0.06 μM was determined (**Figure 6C**). In this case, complete inhibition was observed. The sub micromolar EC_50_ corroborates the high affinity of TP-238 against CECR2.^27^ Next, BZ1 was assessed and demonstrated an EC_50_ of 40 ± 20 μM. BZ1 has the weakest reported affinity for CECR2 and similarly the highest EC_50_ value in our NanoBRET assays. These results indicate that **MS-1** can be an effective tool for determining compound affinities in cells against CECR2.

## CONCLUSIONS

Class I bromodomains are an increasingly disease-relevant class of proteins; however, broadly reactive tracers are lacking and are not commercially available to assess live-cell target engagement and permeability. In this work, we describe the synthesis and assessment of a novel BZ1-based tracer **MS-1**. We show **MS-1** engages with BPTF, PCAF, GCN5, and CECR2 in live cells, and we corroborate literature inhibitor values as well as assess novel in-house compounds for their affinity and permeability. Assessment of this tracer’s compatibility with class IV bromodomains BRD7 and BRD9 would be a future extension of this study. **MS-1** is a valuable tool for the evaluation of novel inhibitors of class I bromo-domains and provides necessary functional in cellulo and permeability data which will bridge the gap between in vitro and in vivo models.

## ASSOCIATED CONTENT

### Supporting Information

The Supporting Information is available free of charge on the ACS Publications website.

Additional NanoBRET dose-response data, Caco-2 permeability methods, HPLC traces of three tracers, general cell culture and NanoBRET assay experimental procedures, small molecule synthetic procedures, and NMR spectra for relevant compounds (PDF).

## AUTHOR INFORMATION

### Author Contributions

M. S. designed and synthesized the NanoBRET tracers; C.T. designed and synthesized the BZ1 derivatives; M.S. performed the cellular experiments; C.T. performed the AlphaScreen assay; X.F. performed the Caco-2 permeability assay; A.A.S. andW.C.K.P oversaw experiments and interpretation of data; M.S. and W.C.K.P wrote the manuscript.

### Notes

No unexpected or unusually high safety hazards were encountered.

The authors declare no competing financial interests.

## ACKNOWLEDGMENT

The authors would like to thank the Promega Corporation for supplying Nluc-fusion vectors for BPTF-BRD, PCAF, CECR2, and GCN5. This work was supported by the National Cancer Institute R01CA290805 and a UMN Muggee Research award. M.S. was additionally supported by a National Institutes of Health Chemical Biology Interface Training Grant T32GM132029. Figures were created with BioRender, ChemDraw, and PyMOL.

## REFERENCES

(1) Filippakopoulos, P.; Picaud, S.; Mangos, M.; Keates, T.; Lambert, J.-P.; Barsyte-Lovejoy, D.; Felletar, I.; Volkmer, R.; Müller, S.; Pawson, T.; Gingras, A.-C.; Arrowsmith, C. H.; Knapp, S. Histone Recognition and Large-Scale Structural Analysis of the Human Bromodomain Family. Cell 2012, 149 (1), 214–231.

(2) Deato, M. D. E.; Tjian, R. Switching of the Core Transcription Machinery during Myogenesis. Genes Dev. 2007, 21 (17), 2137–2149.

(3) Maruyama, T.; Farina, A.; Dey, A.; Cheong, J.; Bermudez, V. P.; Tamura, T.; Sciortino, S.; Shuman, J.; Hurwitz, J.; Ozato, K. A Mammalian Bromodomain Protein, Brd4, Interacts with Replication Factor C and Inhibits Progression to S Phase. Molecular and Cellular Biology 2002, 22 (18), 6509–6520.

(4) Fujisawa, T.; Filippakopoulos, P. Functions of Bromodomain-Containing Proteins and Their Roles in Homeostasis and Cancer. Nat Rev Mol Cell Biol 2017, 18 (4), 246–262.

(5) Liu, L.; Yang, C.; Candelario-Jalil, E. Role of BET Proteins in Inflammation and CNS Diseases. Front. Mol. Biosci. 2021, 8.

(6) Korb, E.; Herre, M.; Zucker-Scharff, I.; Darnell, R. B.; Allis, C. D. BET Protein Brd4 Activates Transcription in Neurons and BET Inhibitor Jq1 Blocks Memory in Mice. Nat Neurosci 2015, 18 (10), 1464–1473.

(7) Hall, M. P.; Unch, J.; Binkowski, B. F.; Valley, M. P.; Butler, B. L.; Wood, M. G.; Otto, P.; Zimmerman, K.; Vidugiris, G.; Machleidt, T.; Robers, M. B.; Benink, H. A.; Eggers, C. T.; Slater, M. R.; Meisenheimer, P. L.; Klaubert, D. H.; Fan, F.; Encell, L. P.; Wood, K. V. Engineered Luciferase Reporter from a Deep Sea Shrimp Utilizing a Novel Imidazopyrazinone Substrate. ACS Chem. Biol. 2012, 7 (11), 1848–1857.

(8) Walker, J. R.; Hall, M. P.; Zimprich, C. A.; Robers, M. B.; Duellman, S. J.; Machleidt, T.; Rodriguez, J.; Zhou, W. Highly Potent Cell-Permeable and Impermeable NanoLuc Luciferase Inhibitors. ACS Chem. Biol. 2017, 12 (4), 1028–1037.

(9) Koblan, L. W.; Buckley, D. L.; Ott, C. J.; Fitzgerald, M. E.; Ember, S. W. J.; Zhu, J.-Y.; Liu, S.; Roberts, J. M.; Remillard, D.; Vittori, S.; Zhang, W.; Schonbrunn, E.; Bradner, J. E. Assessment of Bromodomain Target Engagement by a Series of BI2536 Analogues with Miniaturized BET-BRET. ChemMedChem 2016, 11 (23), 2575–2581.

(10) Mujtaba, S.; He, Y.; Zeng, L.; Farooq, A.; Carlson, J. E.; Ott, M.; Verdin, E.; Zhou, M.-M. Structural Basis of Lysine-Acetylated HIV-1 Tat Recognition by PCAF Bromodomain. Molecular Cell 2002, 9 (3), 575–586. 10.1016/S1097-2765(02)00483-5.

(11) Malatesta, M.; Steinhauer, C.; Mohammad, F.; Pandey, D. P.; Squatrito, M.; Helin, K. Histone Acetyltransferase PCAF Is Required for Hedgehog-Gli-Dependent Transcription and Cancer Cell Proliferation. Cancer Res 2013, 73 (20), 6323–6333.

(12) Zhang, M.; Liu, Z. Z.; Aoshima, K.; Cai, W. L.; Sun, H.; Xu, T.; Zhang, Y.; An, Y.; Chen, J. F.; Chan, L. H.; Aoshima, A.; Lang, S. M.; Tang, Z.; Che, X.; Li, Y.; Rutter, S. J.; Bossuyt, V.; Chen, X.; Morrow, J. S.; Pusztai, L.; Rimm, David. L.; Yin, M.; Yan, Q. CECR2 Drives Breast Cancer Metastasis by Promoting NF-κB Signaling and Macrophage-Mediated Immune Suppression. Sci Transl Med 2022, 14 (630), eabf5473.

(13) Li, B.; Sun, J.; Dong, Z.; Xue, P.; He, X.; Liao, L.; Yuan, L.; Jin, Y. GCN5 Modulates Osteogenic Differentiation of Periodontal Ligament Stem Cells through DKK1 Acetylation in Inflammatory Microenvironment. Sci Rep 2016, 6 (1), 26542..

(14) Peterson, K. E.; Olson, N. M.; Dahlseid, D. J.; Artymiuk, J. M.; Erber, L.; Vitorino, F. N. L.; Dean, R.; Landry, J. W.; Tretyakova, N. Y.; Garcia, B. A.; Pomerantz, W. C. K. BPTF Target Engagement by Acetylated H2A.Z Photoaffinity Probes. Biochemistry 2025, 64 (18), 3872–3885.

(15) Jafari, R.; Almqvist, H.; Axelsson, H.; Ignatushchenko, M.; Lundbäck, T.; Nordlund, P.; Molina, D. M. The Cellular Thermal Shift Assay for Evaluating Drug Target Interactions in Cells. Nat Protoc 2014, 9 (9), 2100–2122.

(16) Picaud, S.; Leonards, K.; Lambert, J.-P.; Dovey, O.; Wells, C.; Fedorov, O.; Monteiro, O.; Fujisawa, T.; Wang, C.-Y.; Lingard, H.; Tallant, C.; Nikbin, N.; Guetzoyan, L.; Ingham, R.; Ley, S. V.; Brennan, P.; Muller, S.; Samsonova, A.; Gingras, A.-C.; Schwaller, J.; Vassiliou, G.; Knapp, S.; Filippakopoulos, P. Promiscuous Targeting of Bromodomains by Bromosporine Identifies BET Proteins as Master Regulators of Primary Transcription Response in Leukemia. Sci. Adv. 2016, 2 (10).

(17) Perell, G. T.; Mishra, N. K.; Sudhamalla, B.; Ycas, P. D.; Islam, K.; Pomerantz, W. C. K. Specific Acetylation Patterns of H2A.Z Form Transient Interactions with the BPTF Bromodomain. Biochemistry 2017, 56 (35), 4607–4615.

(18) Martinelli, P.; Schaaf, O.; Mantoulidis, A.; Martin, L. J.; Fuchs, J. E.; Bader, G.; Gollner, A.; Wolkerstorfer, B.; Rogers, C.; Balıkçı, E.; Lipp, J. J.; Mischerikow, N.; Doebel, S.; Gerstberger, T.; Sommergruber, W.; Huber, K. V. M.; Böttcher, J. Discovery of a Chemical Probe to Study Implications of BPTF Bromodomain Inhibition in Cellular and in Vivo Experiments. ChemMedChem 2023, 18 (6), e202200686.

(19) Zahid, H.; Buchholz, C. R.; Singh, M.; Ciccone, M. F.; Chan, A.; Nithianantham, S.; Shi, K.; Aihara, H.; Fischer, M.; Schönbrunn, E.; dos Santos, C. O.; Landry, J. W.; Pomerantz, W. C. K. New Design Rules for Developing Potent Cell-Active Inhibitors of the Nucleosome Remodeling Factor (NURF) via BPTF Bromodomain Inhibition. J. Med. Chem. 2021, 64 (18), 13902–13917.

(20) Qiang, J.; Zhao, C.; Shi, L.-Q.; Sun, S.-R.; Wang, H.-K.; Liu, S.-L.; Yang, Z.-Y.; Dong, P.; Xiang, S.-S.; Wang, J.-D.; Shu, Y.-J. BRD9 Promotes the Progression of Gallbladder Cancer via CST1 Upregulation and Interaction with FOXP1 through the PI3K/AKT Pathway and Represents a Therapeutic Target. Gene Ther 2024, 31 (11), 594–606.

(21) Yu, X.; Li, Z.; Shen, J. BRD7: A Novel Tumor Suppressor Gene in Different Cancers. Am J Transl Res 2016, 8 (2), 742–748.

(22) Zahid, H.; Costello, J. P.; Li, Y.; Kimbrough, J. R.; Actis, M.; Rankovic, Z.; Yan, Q.; Pomerantz, W. C. K. Design of Class I/IV Bromodomain-Targeting Degraders for Chromatin Remodeling Complexes. ACS Chem. Biol. 2023, 18 (6), 1278–1293.

(23) Lu, T.; Lu, H.; Duan, Z.; Wang, J.; Han, J.; Xiao, S.; Chen, H.; Jiang, H.; Chen, Y.; Yang, F.; Li, Q.; Chen, D.; Lin, J.; Li, B.; Jiang, H.; Chen, K.; Lu, W.; Lin, H.; Luo, C. Discovery of High-Affinity Inhibitors of the BPTF Bromodomain. J Med Chem 2021, 64 (16), 12075–12088.

(24) NanoBRET^TM^ TE 590 Dyes Technical Manual. https://www.promega.com/resources/protocols/technical-manu-als/500/nanobret-te-590-dyes-protocol-tm697/ (accessed 2025-11-15).

(25) Yang, X.; Smith, J. L.; Beck, M. T.; Wilkinson, J. M.; Michaud, A.; Vasta, J. D.; Robers, M. B.; Willson, T. M. Development of Cell Permeable NanoBRET Probes for the Measurement of PLK1 Target Engagement in Live Cells. Molecules 2023, 28 (7), 2950.

(26) Mélin, L.; Calosing, C.; Kharenko, O. A.; Hansen, H. C.; Gagnon, A. Synthesis of NVS-BPTF-1 and Evaluation of Its Biological Activity. Bioorganic & Medicinal Chemistry Letters 2021, 47, 128208.

(27) TP-238 | Structural Genomics Consortium. https://www.thesgc.org/chemical-probes/TP-238 (accessed 2025-10-15).

(28) Urick, A. K.; Hawk, L. M. L.; Cassel, M. K.; Mishra, N. K.; Liu, S.; Adhikari, N.; Zhang, W.; dos Santos, C. O.; Hall, J. L.; Pomerantz, W. C. K. Dual Screening of BPTF and Brd4 Using Protein-Observed Fluorine NMR Uncovers New Bromodomain Probe Molecules. ACS Chem. Biol. 2015, 10 (10), 2246–2256.

(29) Moustakim, M.; Clark, P. G. K.; Trulli, L.; Fuentes de Arriba, A. L.; Ehebauer, M. T.; Chaikuad, A.; Murphy, E. J.; Mendez-Johnson, J.; Daniels, D.; Hou, C.-F. D.; Lin, Y.-H.; Walker, J. R.; Hui, R.; Yang, H.; Dorrell, L.; Rogers, C. M.; Monteiro, O. P.; Fedorov, O.; Huber, K. V. M.; Knapp, S.; Heer, J.; Dixon, D. J.; Brennan, P. E. Discovery of a PCAF Bromodomain Chemical Probe. Angewandte Chemie International Edition 2017, 56 (3), 827–831.

