## Supplemental Information for "NanoBRET Tracer Development for Class I Bromodomain Target Engagement in Live Cells"

**NanoBRET Data** ........ S2
**Caco-2 Cell-Based Permeability Assay** ........ S4
**Analytical HPLC traces of tracers** ........ S5
**NanoBRET Fusion Vectors** ........ S5
**HEK293T Cell Culture and Transfection** ........ S5
**General NanoBRET Assay Protocol** ........ S6
**NanoBRET Tracer Syntheses** ........ S6
**Synthesis of BZ1 Derivatives** ........ S9
**$^{1}$H and $^{13}$C spectra of small molecules** ........ S12
**References** ........ S20

### NanoBRET Data

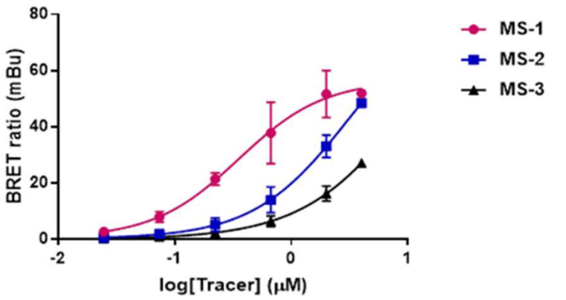


| Tracer | MS-1 | MS-2 | MS-3 |
|---|---|---|---|
| $EC_{50}$ (μM) (n=2) | 0.5 ± 0.1 | 3.2 ± 0.1 | 5.0* |

**Figure S1.** Tracer titration plots against Nluc-BPTF-BRD in HEK293T cells and their $EC_{50}$ values. *second replicate $EC_{50}$ could not be determined due to incomplete curve.

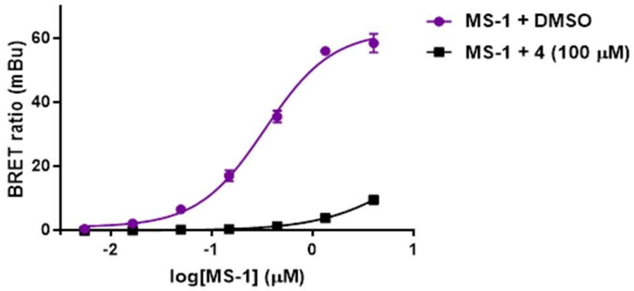


**Figure S2. MS-1** titration against BPTF-BRD-Nluc in the presence of either DMSO (purple) or compound **4** (black) at 100 μM.

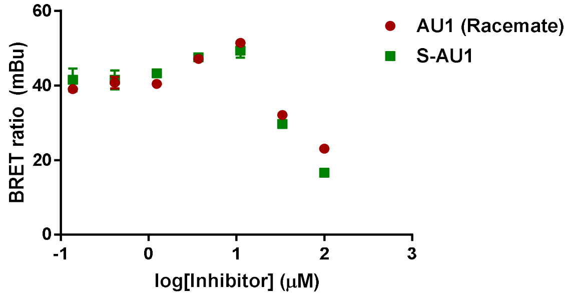

**Figure S3.** Titration plots of AU1 (racemate) and S-AU1 against BPTF-BRD-Nluc using 0.5 μM **MS-1**.

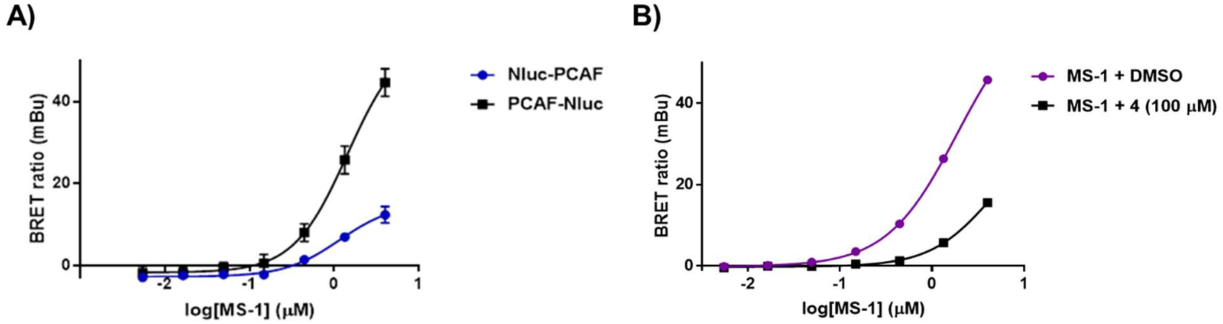


| Construct | Nluc-PCAF | PCAF-Nluc |
| --- | --- | --- |
| MS-1 $EC_{50}$ (μM) (n=1) | 1.2 | 1.4 |

**Figure S4. A) MS-1** titration against Nluc-PCAF (blue) and PCAF-Nluc (black). **B) MS-1** titration against PCAF-Nluc in the presence of either DMSO (purple) or compound **4** (black) at 100 μM.

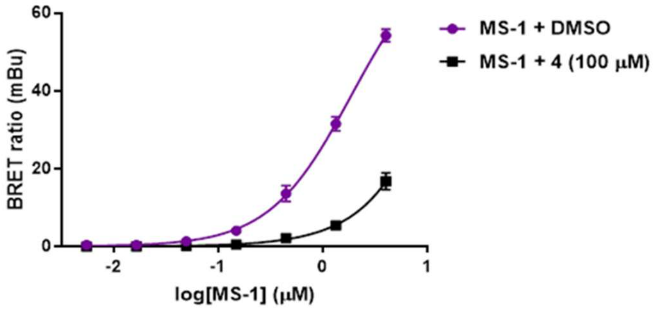


| Construct | Nluc-GCN5 |
| --- | --- |
| MS-1 $EC_{50}$ (μM) (n=1) | 1.9 |

**Figure S5. MS-1** titration against Nluc-GCN5 in the presence of either DMSO (purple) or compound **4** (black) at 100 μM.

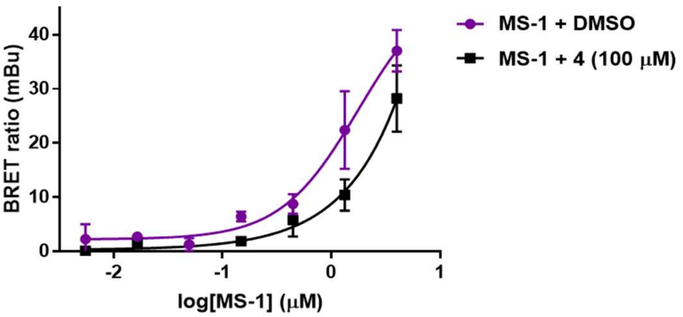


| Construct | Nluc-CECR2 |
| --- | --- |
| MS-1 $EC_{50}$ (μM) (n=1) | 1.8 |

**Figure S6. MS-1** titration against Nluc-CECR2 in the presence of either DMSO (purple) or compound **4** (black) at 100 µM.

### Caco-2 Cell-Based Permeability Assay

The Caco-2 permeability assay was conducted as previously described (PMID: 36793425). Briefly, Caco-2 cells were maintained at 37 °C in a humidified incubator with 5% CO2, cultured in MEM supplemented with 10% FBS, 100 units/ml of penicillin, and 100 µg/ml of streptomycin in 75 $cm^2$ flasks. Cells were seeded onto 96-well Transwell inserts at a density of $0.165 \times 10^5$ cells/insert and cultured for an additional 7 days. Monolayers were washed twice with HBSS/HEPES (10 mM, pH 7.4) before starting the permeability assay. To initiate the assay, a 10 µmol/L solution of each test compound was added to either the apical (A) or basolateral (B) side of the inserts. The monolayers were incubated for 2 h at 37 °C. Samples were collected from both compartments and compound concentrations were measured by UPLC/MS (Waters; Milford, MA). All compounds were tested in triplicates.

The apparent permeability coefficient (Papp, cm/s) of each compound in the A→B (or B→A) direction was calculated using the equation:

$$Papp = (dQ/dt) \times (1/AC_0),$$

where dQ/dt is the flux (µmol/s), A is the surface area ($cm^2$), and $C_0$ is the initial concentration (µmol/L).

Chromatographic separation was performed on a Waters Acquity UPLC system coupled to an SQ mass spectrometer. Data acquisition used Masslynx v4.1 and analysis was performed with Quanlynx. The flow rate was 1.0 mL/min and the injection volume was 10 µL. The UPLC column (Acquity BEH C18, 1.7 µm, 2.1 × 50 mm) was maintained at 63 °C. The mobile phases were solvent A (0.1% formic acid in MilliQ $H_2O$) and solvent B (0.1% formic acid in acetonitrile), with the following gradient:

- 0–0.2 min: B% 10–30%
- 0.2–1.6 min: B% 30–95%
- 1.6–1.95 min: B% 95%
- 1.95–2 min: B% 95–10%

The mass spectrometer operated in positive-ion electrospray ionization mode with the following settings: capillary voltage 3.4 kV, cone voltage 50 V, source temperature 150 °C, desolvation temperature 350 °C, desolvation gas 800 L/hr, cone gas 25 L/hr. MS data were acquired with a full scan range of m/z 150–1200 (0.2 s scan time). Single ion recording was used for quantification of each compound.

### Analytical HPLC traces of tracers

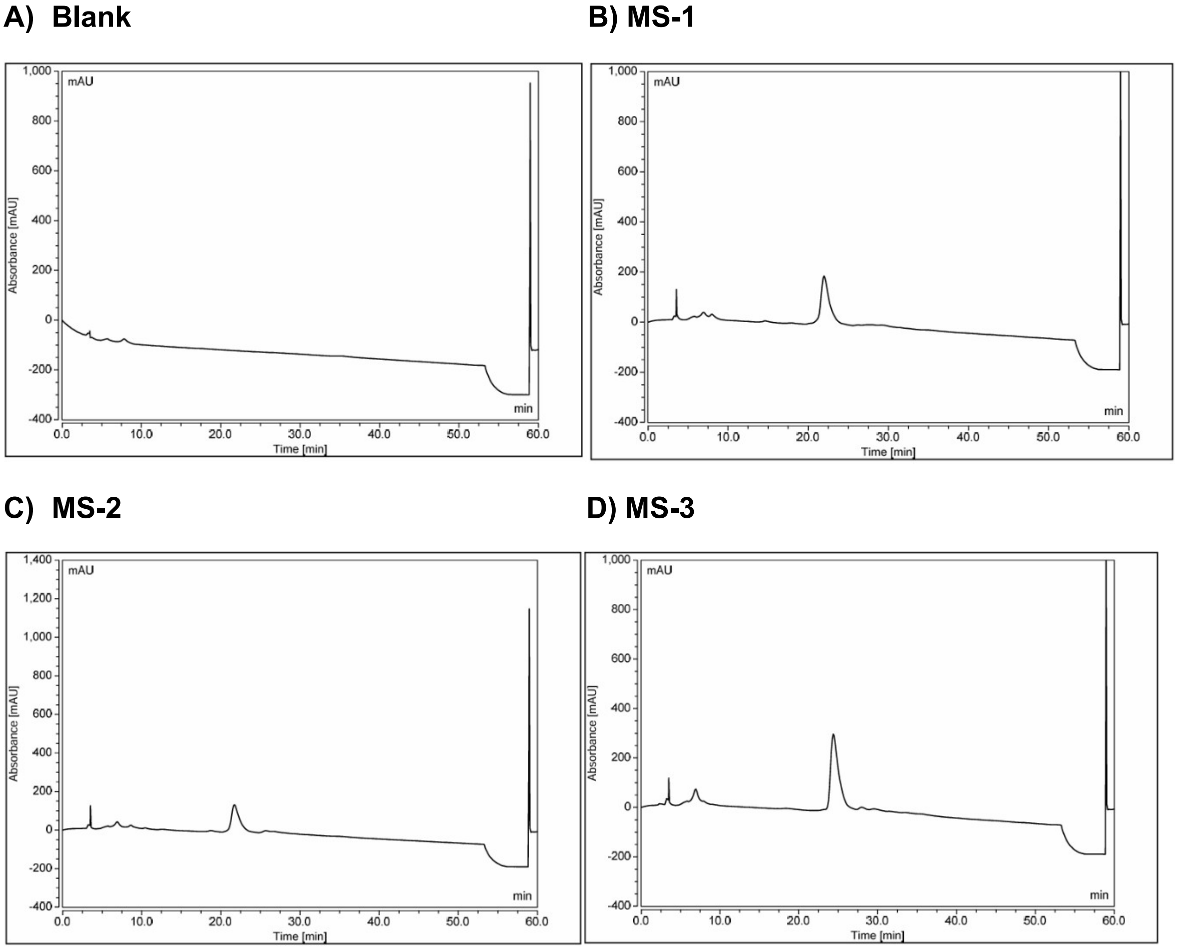


**Figure S7.** HPLC traces at 220 nm of **A)** Blank, **B)** MS-1, **C)** MS-2, **D)** MS-3 over a gradient of 0-50% acetonitrile in 0.1% TFA in water.

### NanoBRET Fusion Vectors

The BPTF-BRD-NanoLuc fusion vector was designed based on previously reported literature.[1] The N- and C-terminal-NanoLuc fusion vectors for PCAF, CECR2, and GCN5 were provided by Promega.

### HEK293T Cell Culture and Transfection

HEK293T cells were cultured in DMEM, 10% FBS, 5% penicillin-streptomycin and incubated at 37 °C with 5% $CO_2$. The NanoBRET assay was executed based on the Promega NanoBRET TE 590 Dyes technical protocol. HEK293T cells (200,000 cells/mL) were transiently transfected with each target-NanoLuc fusion vector. A 10 µg/mL solution of lipid:DNA complexes containing each fusion vector were prepared with 9.0 µg/mL of transfection carrier DNA (Promega), 1.0 µg/mL of target-NanoLuc fusion vector DNA

(Promega), and 1 mL of Opti-MEM I. FuGENE HD Transfection Reagent (Promega) was added at a concentration of 30 µL per mL of DNA mixture. The lipid:DNA complex mixture was inverted five times and allowed to incubate for 20 minutes at room temperature. Lipid:DNA complex was then added to HEK293T cells in a 1:20 lipid:DNA complex to cells ratio. The subsequent mixture was inverted and dispensed into sterile tissue culture flasks at a density between 55,000 and 80,000 cells/$cm^2$. The cells were incubated for 20 – 30 hours at 37 °C, 5% $CO_2$. Following incubation, the transfected cells were then adjusted to a density of 200,000 cells/mL in Opti-MEM I.

#### General NanoBRET Assay Protocol

BRET assays were performed using white, 384-well tissue-culture-treated plates (Corning #3570). HEK293T cells were plated at 7 x $10^3$ cells per well. Stock DMSO solutions of each novel BRET tracer to be tested were diluted in Tracer Dilution Buffer (Promega) to achieve a 20X concentration by adding 4:1 Tracer Dilution Buffer to 1 part 100X tracer. BPTF-BRD BRET assays were conducted using a final **MS-1** concentration of 0.5 µM, and PCAF, GCN5, and CECR2 used a final **MS-1** concentration of 1 µM. Stock DMSO solutions of inhibitors were diluted to a 1,000X concentration in 100% DMSO and subsequently dissolved to a final 10X concentration in Opti-MEM. Upon addition of the tracer and inhibitor, the cells were incubated for 2 hours at 37 °C, 5% $CO_2$. Following incubation, a 3X stock solution of NanoBRET Nano-Glo Substrate and Extracellular NanoLuc Inhibitor in Opti-MEM was dispensed into each well. BRET was then measured in a Tecan Spark plate reader following a 30 second orbital shake with an amplitude of 1 mm and frequency of 510 rpm. Luminescence was measured using the luminescence multi color setting with a donor emission of 445 nm – 470 nm and acceptor emission of 610 nm – 700 nm using an integration time of 300 ms. Each experiment also included DMSO control wells with no tracer and no inhibitor for background correction as well as control wells with tracer and no inhibitor to determine the maximal assay window. For both tracer and inhibitor titration plots, a BRET ratio for each concentration was calculated as acceptor response divided by donor response. The BRET ratio was then subtracted by the no tracer DMSO control well ratio and multiplied by 1000 to obtain a milliBRET (mBRET) value. The mBRET values were plotted and fit using the 4-parameter dose-response variable slope curve in GraphPad Prism.

#### NanoBRET Tracer Syntheses

*tert-butyl (4-(6-((5-chloro-1-methyl-6-oxo-1,6-dihydropyridazin-4-yl)amino)-3,4-dihydroisoquinolin-2(1H)-yl)butyl)carbamate (**2**)*

Compound **1** was synthesized based on previously reported literature.[2] **1** (0.200 g, 1.0 eq, 0.69 mmol) was stirred in 1,4-dioxane (4.2 mL, 163 mM), then 4-(Bocamino)butyl bromide (0.191 g, 1.1 eq, 0.76 mmol) and *N,N*-diisopropylethylamine (0.18 mL, 1.5 eq,

1.0 mmol) were added. The mixture was heated in a sealed tube and stirred at 110 °C for 18 h. Upon completion of the reaction, the 1,4-dioxane was removed by rotary evaporation. Subsequently, the crude mixture extracted into ethyl acetate (3 x 20 mL), washed with saturated sodium bicarbonate (3 x 20 mL), and with brine (1 x 20 mL). The organic layer was dried with magnesium sulfate, then concentrated by rotary evaporation and purified by flash chromatography to yield compound **2** as a white solid (45% yield). (CombiFlash Rf system: 4 g of silica, DCM/methanol, 0-20% methanol, 30 min). $^1$H NMR (500 MHz, chloroform-*d*) δ 7.61 (s, 1H), 7.65 (d, *J* = 8.0 Hz, 1H), 6.96 (s, 1H), 6.95 (d, *J* = 7.6 Hz, 1H), 6.41 (s, 1H), 5.00 (br. s, 1H), 3.74 (s, 3H), 3.68 (s, 2H), 3.15 (q, *J* = 6.5 Hz, 2H), 2.94 (t, *J* = 5.9 Hz, 2H), 2.79 (t, *J* = 6.0 Hz, 2H), 2.60 (t, *J* = 7.4 Hz, 2H), 1.67 (quintet, *J* = 7.4 Hz, 2H), 1.57 (quintet, *J* = 7.2 Hz, 2H), 1.41 (s, 9H), DCM impurity at 5.29 ppm (4.8% by weight). $^{13}$C NMR (126 MHz, chloroform-*d*) δ 157.8, 156.1, 142.2, 136.0, 135.5, 128.1, 126.7, 124.2, 121.9, 109.6, 79.0, 77.3, 77.1, 76.8, 57.6, 55.3, 50.4, 40.3, 28.7, 28.4, 27.9, 24.1 (two resonances overlapping). HRMS (ESI-TOF) calculated for $C_{23}H_{33}ClN_5O_3^+$ [M+H]$^+$ = 462.2271, observed 462.2242.

*5-((2-(4-aminobutyl)-1,2,3,4-tetrahydroisoquinolin-6-yl)amino)-4-chloro-2-methylpyridazin-3(2H)-one (**3**)*

Compound **2** (0.081 g, 1.0 eq, 0.175 mmol) was dissolved in dichloromethane (1 mL, 175 mM) and stirred at room temperature with trifluoroacetic acid (0.067 mL, 5.0 eq, 0.876 mmol) for 18 h. Upon completion of the reaction, the dichloromethane and residual TFA was removed using an $N_2$ stream to yield compound **3** as a brown oil (4x TFA salt, quantitative yield). $^1$H NMR (500 MHz, DMSO-*d$_6$*) δ 8.80 (s, 1H), 7.94 (br. s, 3H), 7.66 (s, 1H), 7.22 (d, *J* = 8.3 Hz, 1H), 7.16 (dd, *J* = 8.2, 2.3 Hz, 1H), 7.14 (d, *J* = 2.2 Hz, 1H), 4.54 (d, *J* = 15.4 Hz, 1H), 4.27 (dd, J = 15.9, 6.6 Hz, 1 H), 3.75 – 3.67 (m, 1H), 3.62 (s, 3H), 3.42 – 3.00 (m, 5H), 2.86 (sextet, *J* = 6.5 Hz, 2 H) 1.81 (quintet, *J* = 7.9 Hz, 2H), 1.62 (quintet, *J* = 7.6 Hz, 2H), TFA impurity at 10.4 ppm (23% by weight). $^{13}$C NMR (126 MHz, DMSO-*d$_6$*) δ 158.4, 157.1, 142.1, 138.1, 132.5, 127.8, 127.7, 124.6, 122.7, 121.8, 109.2, 54.3, 51.7, 48.6, 38.2, 24.8, 24.2, 20.6 (two resonances overlapping). HRMS (ESI-TOF) calculated for $C_{18}H_{25}ClN_5O^+$ [M+H]$^+$ = 362.1748, observed 362.1743.

General procedure for synthesis of three tracers

Compound **3** was dissolved in dry dimethyl formamide (1 mL, 8.1 M) and stirred at room temperature with *N,N*-diisopropylethylamine (30.6 µL, 15 eq, 176 mmol) in an amber vial. Then, the BODIPY578/589 (5 mg, 1.2 eq) with either an alkyl or PEG linker construct was added and the vial was covered in aluminum foil to protect the fluorophore from light. The mixture was allowed to stir overnight. Upon reaction completion, the mixture was extracted into dichloromethane (3 x 20 mL), then washed with deionized water (1 x 20 mL) and once with brine (1 x 20 mL). The organic layer was dried with magnesium sulfate, then concentrated by rotary evaporation and purified by flash chromatography to yield the final compound as a purple solid. (Combiflash Rf system: 4 g of silica, DCM/methanol, 0-100% methanol, 30 min).

*N-(4-(6-((5-chloro-1-methyl-6-oxo-1,6-dihydropyridazin-4-yl)amino)-3,4-dihydroisoquinolin-2(1H)-yl)butyl)-3-(5,5-difluoro-7-(1H-pyrrol-2-yl)-5H-5l4,6l4-dipyrrolo[1,2-c:2',1'-f][1,3,2]diazaborinin-3-yl)propenamide (**MS-1**)*

**MS-1** was synthesized following general procedure A with 21.4 mg compound **3** (1.0 eq, 9.8 µmol) and obtained as a purple solid (quantitative yield). HRMS (ESI-TOF) calculated for $C34H37BClF2N8O2^{+}$ $[M+H]^{+}$ = 673.2789, observed 673.2620.

*N-(4-(6-((5-chloro-1-methyl-6-oxo-1,6-dihydropyridazin-4-yl)amino)-3,4-dihydroisoquinolin-2(1H)-yl)butyl)-4-(3-(5,5-difluoro-7-(1H-pyrrol-2-yl)-5H-5l4,6l4-dipyrrolo[1,2-c:2',1'-f][1,3,2]diazaborinin-3-yl)propanamido)butanamide (**MS-2**)*

**MS-2** was synthesized following general procedure A with 17.8 mg compound **3** (1.0 eq, 8.1 µmol) and obtained as a purple solid (96% yield). HRMS (ESI-TOF) calculated for $C38H44BClF2N9O3^{+}$ $[M+H]^{+}$ = 758.3317, observed 758.3094.

*N-(4-(6-((5-chloro-1-methyl-6-oxo-1,6-dihydropyridazin-4-yl)amino)-3,4-dihydroisoquinolin-2(1H)-yl)butyl)-1-(3-(5,5-difluoro-7-(1H-pyrrol-2-yl)-5H-5l4,6l4-dipyrrolo[1,2-c:2',1'-f][1,3,2]diazaborinin-3-yl)propanamido)-3,6,9,12-tetraoxapentadecan-15-amide (**MS-3**)*

**MS-3** was synthesized following general procedure A with 13.6 mg compound **3** (1.0 eq, 6.2 µmol) and obtained as a purple solid (68% yield). HRMS (ESI-TOF) calculated for $C45H58BClF2N9O7^{+}$ $[M+H]^{+}$ = 920.4209, observed 920.4443.

### Synthesis of BZ1 Derivatives

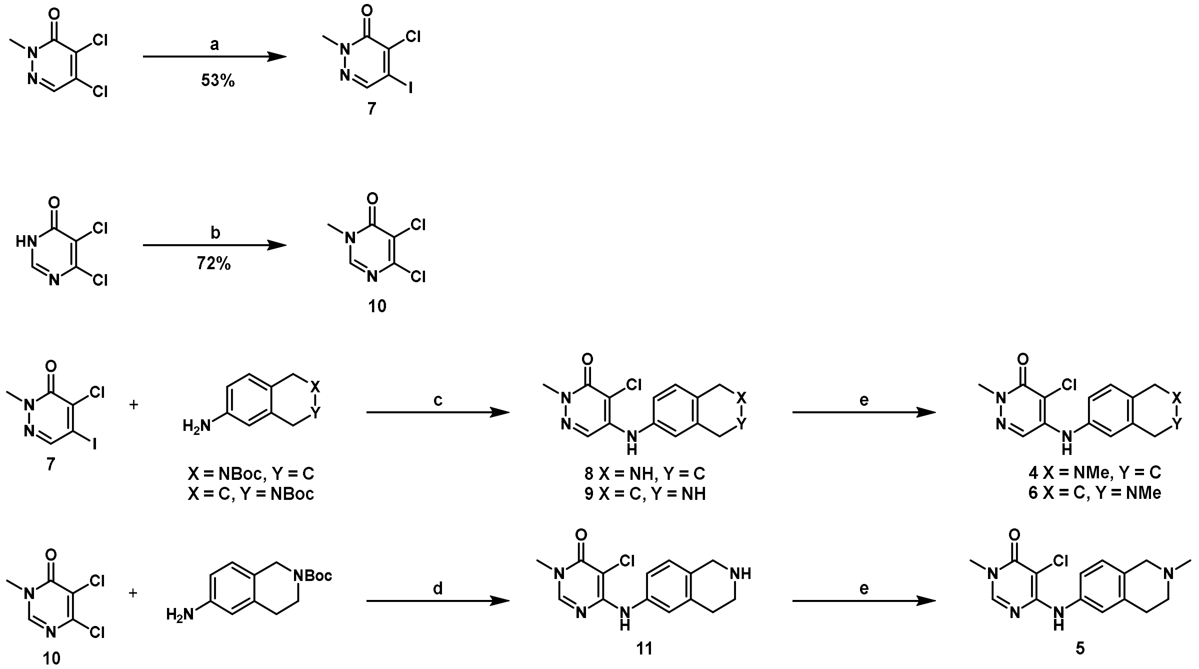


*4-chloro-5-iodo-2-methylpyridazin-3(2H)-one* ***(7)***

4,5-dichloro-2-methylpyridazin-3(2H)-one (30 g, 0.17 mol, 1 eq), sodium iodide (75 g, 0.5 mol, 1.2 eq) were added to DMF (335 mL). The reaction mixture was stirred at 180 °C for 24 hours and was washed with 20% lithium chloride solution. The resultant mixture was extracted with ethyl acetate and concentrated in vacuo. The crude was then dissolved in a minimal amount of boiling methanol and cooled to 4 °C overnight for recrystallization. The crystals were filtered and washed with ether, yielding 4-chloro-5-iodo-2-methylpyridazin-3(2H)-one (53%) as an off-white powder (with a ratio of 6.5:1 compared with the starting material). The filtrate can be concentrated and subjected to recrystallization 2~3 times to get more product. $^{1}H$ NMR (500 MHz, $CDCl_3$) δ 7.96 (s, 1H), 3.78 (s, 3H); $^{13}C$ NMR (126 MHz, $CDCl_3$) δ 155.89, 142.33, 141.65, 104.55, 41.00; HRMS (ESI) calculated for $C_6H_5ClINO$ $[M + H]^+$, 270.9131; observed, 270.9153.

*5,6-dichloro-3-methylpyrimidin-4(3H)-one* ***(10)***

5,6-dichloropyrimidin-4(3*H*)-one (445 mg, 2.70 mmol, 1 eq), methyl iodide (460 mg, 3.24 mmol, 1.2 eq), and potassium carbonate (447 mg, 3.23 mmol, 1.2 eq) were added to THF (2.7 mL). The reaction mixture was stirred at room temperature overnight and was washed with 10% sodium thiosulfate solution. The resultant mixture was extracted with ethyl acetate (3 × 20 mL), and the filtrate was concentrated in vacuo, yielding 5,6-dichloro-3-methylpyrimidin-4(3*H*)-one (72%) as a crude. $^{1}H$ NMR (500 MHz, $CDCl_3$) δ 7.98 (s, 1H), 3.58 (s, 3H); $^{13}C$ NMR (126 MHz, $CDCl_3$) δ 171.27, 154.66, 147.43, 121.31, 35.27; HRMS (ESI) calculated for $C_5H_4Cl_2N_2O$ $[M + H]^+$, 178.9773; observed, 178.9796.

*4-chloro-2-methyl-5-((1,2,3,4-tetrahydroisoquinolin-6-yl)amino)pyridazin-3(2H)-one **(8)***

To a solution of 4-chloro-5-iodo-2-methylpyridazin-3(2H)-one **(7)** (1 g, 3.69 mmol, 1 eq) in dry toluene (18 mL) was added tert-butyl 6-amino-3,4-dihydroisoquinoline-2(1H)-carboxylate (1.4 g, 5.54 mmol, 1.5 eq), cesium carbonate (2.9 mg, 8.87 mmol, 2.4 eq) and a freshly prepared solution of $Pd(OAc)_2$/Rac-BINAP in dry toluene. This solution was obtained by stirring $Pd(OAc)_2$ (50 mg, 0.22 mmol, 0.06 eq) and Rac-BINAP (207 mg, 0.33 mmol, 0.09 eq) in dry toluene (18 mL) for 30 minutes with argon bubbling through the mixture. The main reaction mixture was heated at 110 °C for 16 h, cooled down to room temperature, filtered over silica gel, rinsed with a mixture of ethyl acetate, and concentrated under reduced pressure and purified by flash column chromatography to yield tert-butyl 6-((5-chloro-1-methyl-6-oxo-1,6-dihydropyridazin-4-yl)amino)-3,4-dihydroisoquinoline-2(1H)-carboxylate (1 g,73% yield). To remove the Boc protecting group, the yielded product was stirred in DCM, and to the mixture was added trifluoroacetic acid (10.0 eq)) at room temperature overnight. The reaction was then extracted into DCM before being treated with 1 M NaOH to achieve a pH of 10. The DCM layer was then dried over magnesium sulfate, filtered, and purified by flash column chromatography to yield the product **8** (500 mg, 47% overall yield). $^1$H NMR (500 MHz, MeOD) δ 7.72 (s, 1H), 7.29 (d, $J$ = 8.2 Hz, 1H), 7.22 – 7.15 (m, 2H), 4.37 (s, 2H), 3.73 (s, 3H), 3.51 (t, $J$ = 6.4 Hz, 2H), 3.14 (t, $J$ = 6.4 Hz, 2H). $^{13}$C NMR (126 MHz, MeOD) δ 158.71, 143.26, 137.98, 133.13, 127.95, 127.91, 125.57, 124.08, 122.79, 108.83, 44.16, 41.24, 39.30, 24.73. HRMS (ESI) calculated for $C_{14}H_{15}ClN_4O$ $[M + H]^+$, 291.1007; observed, 291.0996.

*4-chloro-2-methyl-5-((2-methyl-1,2,3,4-tetrahydroisoquinolin-6-yl)amino)pyridazin-3(2H)-one **(4).***

Formaldehyde (37%) in water with 10−15% MeOH (0.26 mL, 3.44 mmol, 10 eq) and formic acid (1 mL) were added to 4-chloro-2-methyl-5-((1,2,3,4-tetrahydroisoquinolin-6-yl)amino)pyridazin-3(2H)-one **(8)** (100 mg, 0.34 mmol, 1 eq) in a sealed tube or refluxed at 80 °C for 6 h. The reaction mixture was allowed to cool down to room temperature and purified by flash column chromatography to yield **4** (115 mg, 100%). $^1$H NMR (400 MHz, $CDCl_3$) δ 7.67 (s, 1H), 7.13 (d, $J$ = 8.2 Hz, 1H), 7.09 – 6.98 (m, 2H), 6.38 (s, 1H), 4.04 (s, 2H), 3.78 (s, 3H), 3.18 (s, 4H), 2.78 (s, 3H); $^{13}$C NMR (126 MHz, $CDCl_3$) δ 157.91, 142.02, 136.56, 134.47, 128.22, 126.78, 124.02, 122.23, 56.10, 51.92, 44.73, 40.46, 27.64; HRMS (ESI) calculated for $C_{15}H_{17}ClN_4O$ $[M + H]^+$, 305.1164; observed, 305.1156.

*5-chloro-3-methyl-6-((1,2,3,4-tetrahydroisoquinolin-6-yl)amino)pyrimidin-4(3H)-one **(11)***

5,6-dichloro-3-methylpyrimidin-4(3*H*)-one (200 mg, 1.12 mmol, 1 eq), tert-butyl (2-(4-amino-phenyl)-ethyl)carbamate (248 mg, 1 mmol, 0.9 eq), and N, N-diisopropylethylamine (288 mg, 2.24 mmol, 2.0 eq) were dissolved in DMSO (500 µL) and heated at 120 °C for 16 h. The reaction mixture was extracted with ethyl acetate and washed three times with a 1 M sodium bicarbonate solution, followed by a brine wash. The organic layer was then dried with magnesium sulfate, filtered, concentrated in vacuo, and purified by flash column chromatography to yield tert-butyl 6-((5-chloro-1-methyl-6-oxo-1,6-dihydropyrimidin-4-yl)amino)-3,4-dihydroisoquinoline-2(1H)-carboxylate (215 mg, 49% yield). Both 4- and 5-positional isomers were obtained, with the desired compound being the major product. To remove the Boc protecting group, the yielded product was stirred in DCM (1 mL) at room temperature, and trifluoroacetic acid (10.0 eq) was added to the mixture. The reaction was stirred at room temperature overnight and then extracted into DCM before being treated with 1 M NaOH to adjust the pH to 10. The DCM layer was then dried over magnesium sulfate, filtered, and purified by flash column chromatography to yield compound **11**.

*5-chloro-3-methyl-6-((2-methyl-1,2,3,4-tetrahydroisoquinolin-6-yl)amino)pyrimidin-4(3H)-one* ***(5).***

Formaldehyde (37%) in water with 10−15% MeOH (0.45 mL, 5.51 mmol, 10 eq) and formic acid (1.6 mL) were added to 5-chloro-3-methyl-6-((1,2,3,4-tetrahydroisoquinolin-6-yl)amino)pyrimidin-4(3H)-one **(11)** (167 mg, 0.55 mmol, 1 eq) in a sealed tube or refluxed at 80 °C for 6 h. The reaction mixture was allowed to cool down to room temperature and purified by flash column chromatography to yield **5** (130 mg, 77%). $^{1}H$ NMR (400 MHz, MeOD) δ 8.09 (s, 1H), 7.45 – 7.36 (m, 2H), 7.17 – 7.07 (m, 1H), 4.41 (s, 2H), 3.58 (s, 2H), 3.47 (s, 3H), 3.26 – 3.13 (m, 2H), 3.05 (s, 3H); $^{13}C$ NMR (126 MHz, $CDCl_3$) δ 158.49, 155.07, 148.09, 138.23, 130.99, 127.46, 122.59, 121.90, 121.43, 97.57, 54.10, 50.85, 41.95, 34.70, 24.91; HRMS (ESI) calculated for $C_{15}H_{17}ClN_4O$ $[M + H]^+$, 305.1164; observed, 305.1169.

*4-chloro-2-methyl-5-((1,2,3,4-tetrahydroisoquinolin-7-yl)amino)pyridazin-3(2H)-one* ***(9)***

To a solution of 4-chloro-5-iodo-2-methylpyridazin-3(2H)-one **(7)** (500 mg, 1.85 mmol, 1 eq) in dry toluene (9.3 mL) was added tert-butyl 7-amino-3,4-dihydroisoquinoline-2(1H)-carboxylate (689 mg, 2.77 mmol, 1.5 eq), cesium carbonate (1.45 g, 4.44 mmol, 2.4 eq) and a freshly prepared solution of $Pd(OAc)_2$/Rac-BINAP in dry toluene. This solution was obtained by stirring $Pd(OAc)_2$ (25 mg, 0.11 mmol, 0.06 eq) and Rac-BINAP (103 mg, 0.17 mmol, 0.09 eq) in dry toluene (9.3 mL) for 30 min with argon bubbling through the mixture. The main reaction mixture was heated at 110 °C for 16 h, cooled down to room temperature, filtered over silica gel, rinsed with a mixture of ethyl acetate, and concentrated under reduced pressure and purified by flash column chromatography to yield tert-butyl 7-((5-chloro-1-methyl-6-oxo-1,6-dihydropyridazin-4-yl)amino)-3,4-dihydroisoquinoline-2(1H)-carboxylate (490 mg, 68% yield). To remove the Boc protecting group, the yielded product was stirred in DCM (5 mL) at room temperature, and trifluoroacetic acid (10.0 eq) was added to the mixture. The reaction was stirred at room temperature overnight and then extracted into DCM before being treated with 1 M NaOH to adjust the pH to 10. The DCM layer was then dried over magnesium sulfate, filtered, and removed under vacuum to obtain product **9** as a TFA salt.

*4-chloro-2-methyl-5-((2-methyl-1,2,3,4-tetrahydroisoquinolin-7-yl)amino)pyridazin-3(2H)-one **(6)***

Formaldehyde (37%) in water with 10−15% MeOH (0.26 mL, 3.44 mmol, 10 eq) and formic acid (1 mL) were added to 4-chloro-2-methyl-5-((1,2,3,4-tetrahydroisoquinolin-7-yl)amino)pyridazin-3(2H)-one **(9)** (100 mg, 0.34 mmol, 1 eq) in a sealed tube or refluxed at 80 °C for 6 h. The reaction mixture was allowed to cool down to room temperature and purified by flash column chromatography to yield **6** (60 mg, 58%). $^1$H NMR (500 MHz, MeOD) δ 7.71 (s, 1H), 7.32 (d, *J* = 8.3 Hz, 1H), 7.20 (dd, *J* = 8.2, 2.3 Hz, 1H), 7.11 (d, *J* = 2.4 Hz, 1H), 4.39 (s, 2H), 3.71 (s, 3H), 3.54 (t, *J* = 6.4 Hz, 2H), 3.20 (t, *J* = 6.3 Hz, 2H), 3.01 (s, 3H); $^{13}$C NMR (126 MHz, CDCl3) δ 157.90, 141.79, 136.72, 130.70, 130.16, 128.92, 126.61, 123.80, 121.80, 110.84, 54.54, 51.17, 42.87, 40.53, 25.29; HRMS (ESI) calculated for $C_{15}H_{17}ClN_4O$ [M + H]$^+$, 305.1164; observed, 305.1136.

**$^1$H and $^{13}$C spectra of small molecules**

**2**, $^1$H NMR (500 MHz, chloroform-*d*)

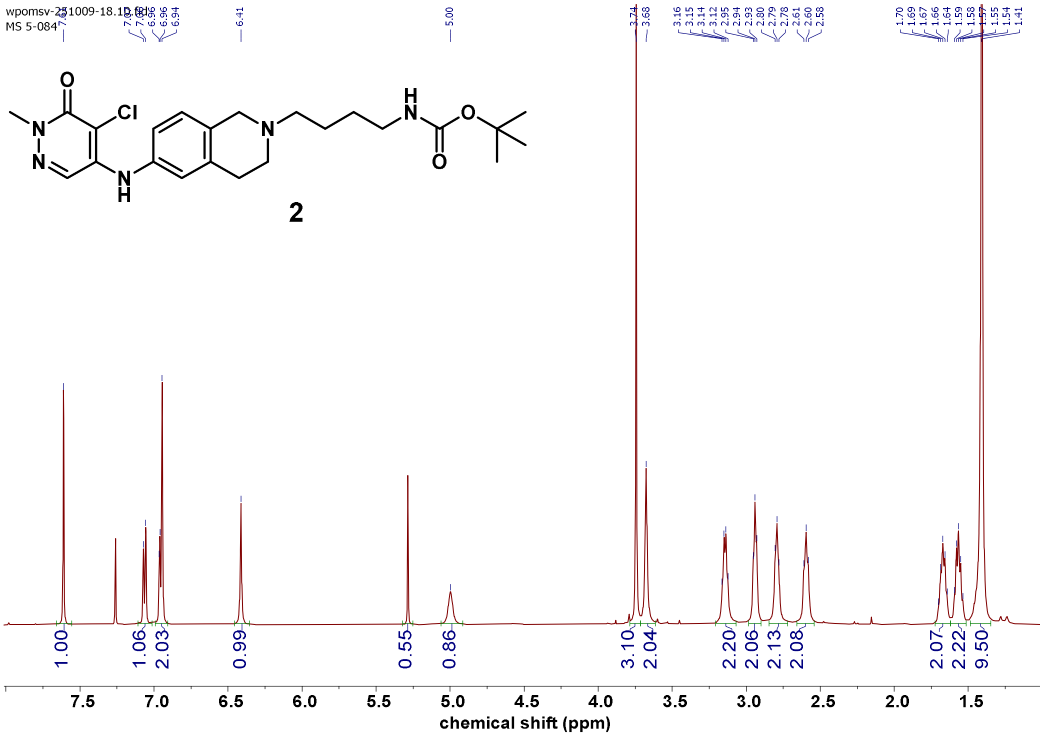


**2**, $^{13}$C NMR (126 MHz, chloroform-*d*)

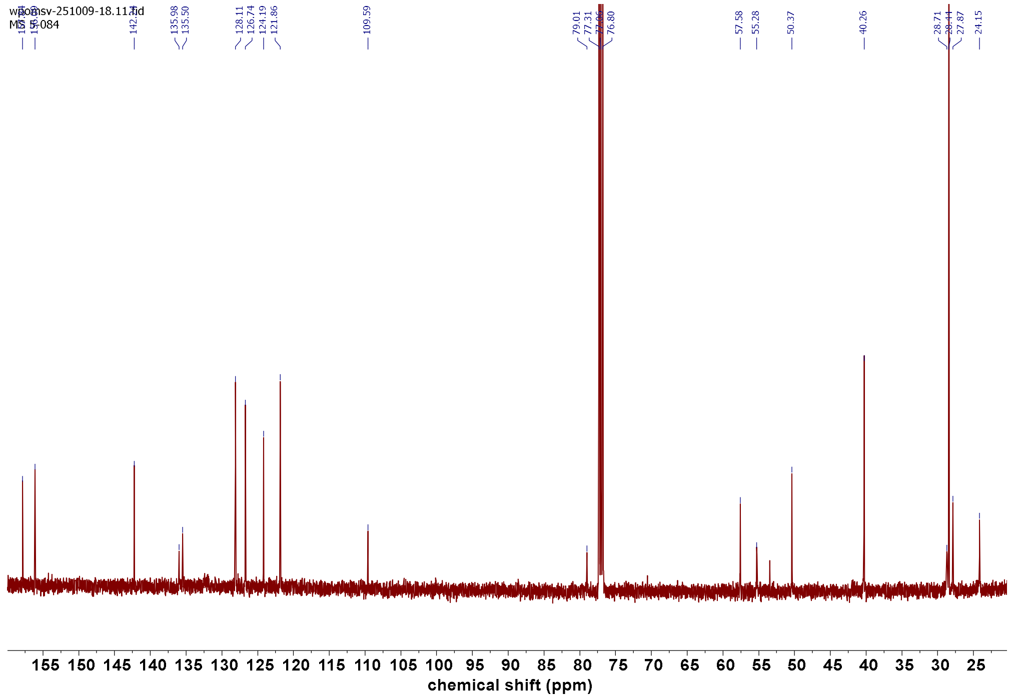


3, ¹H NMR (500 MHz, DMSO-$d_6$)

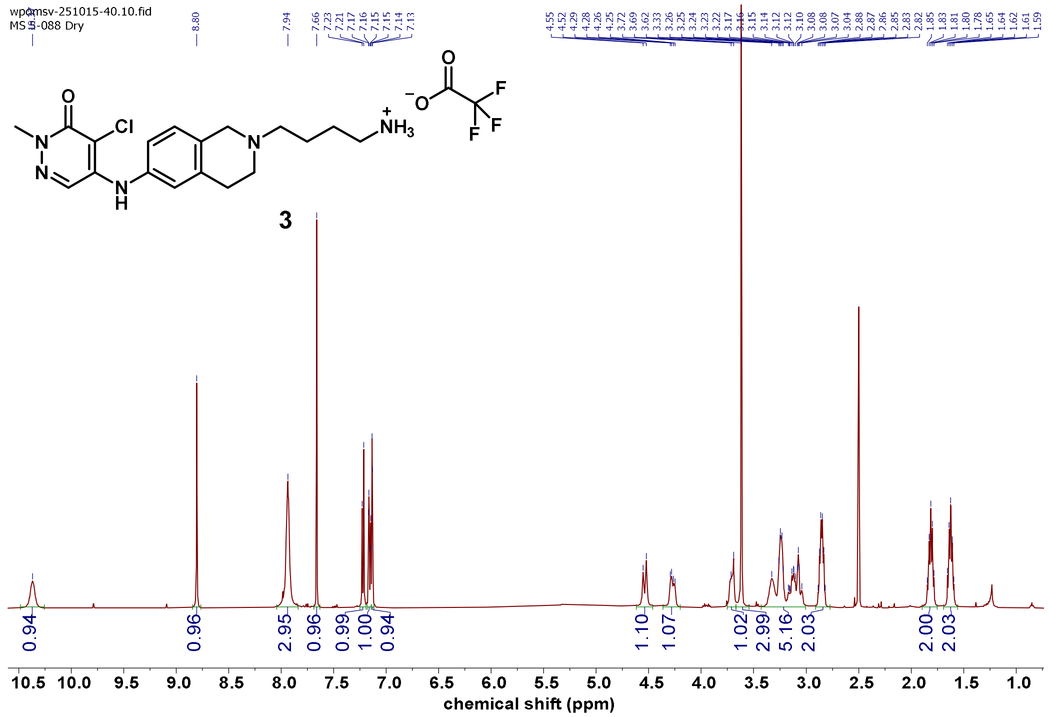


3, ¹³C NMR (126 MHz, DMSO-$d_6$)

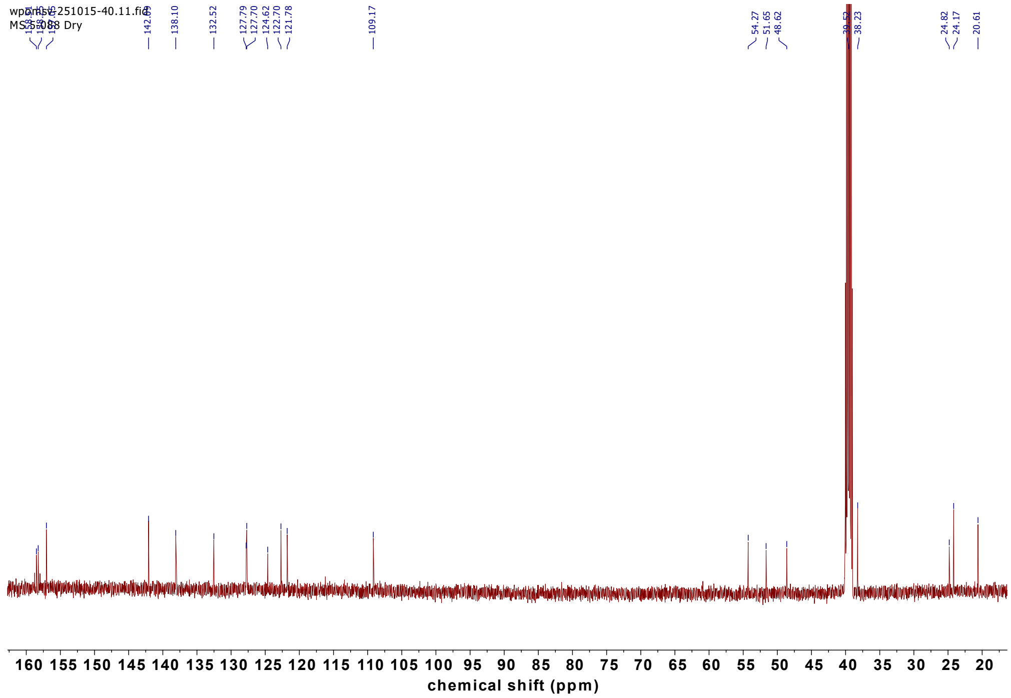

**7**, $^{1}$H NMR (500 MHz, Chloroform-*d*)

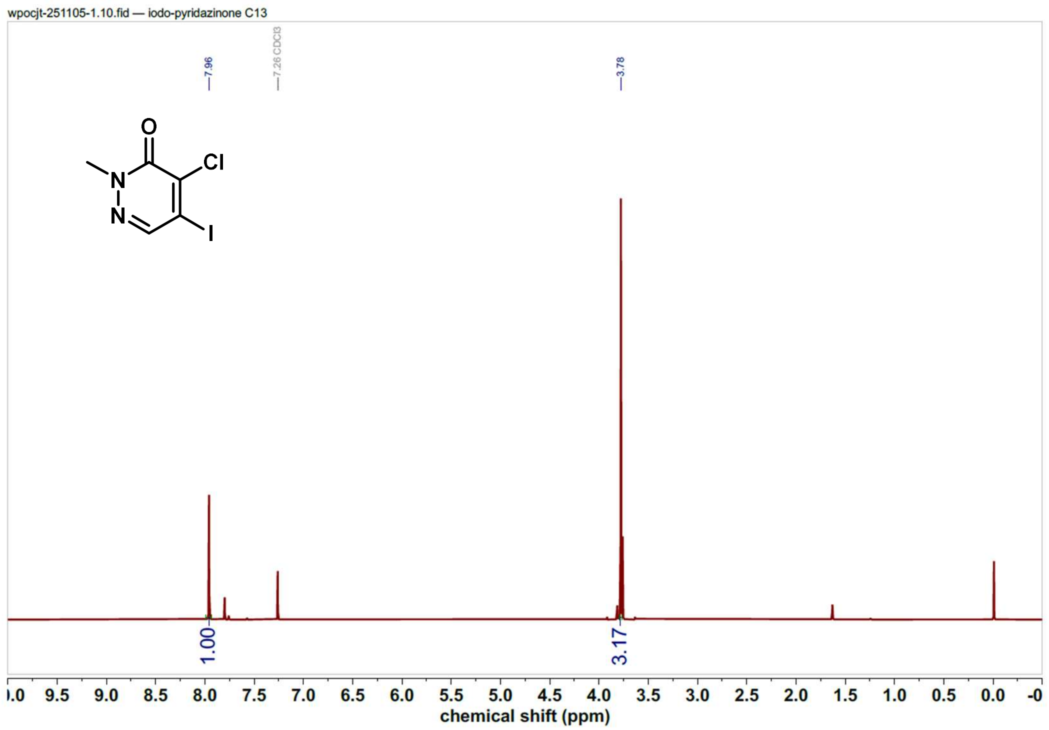

**7**, $^{13}$C NMR (126 MHz, Chloroform-*d*)

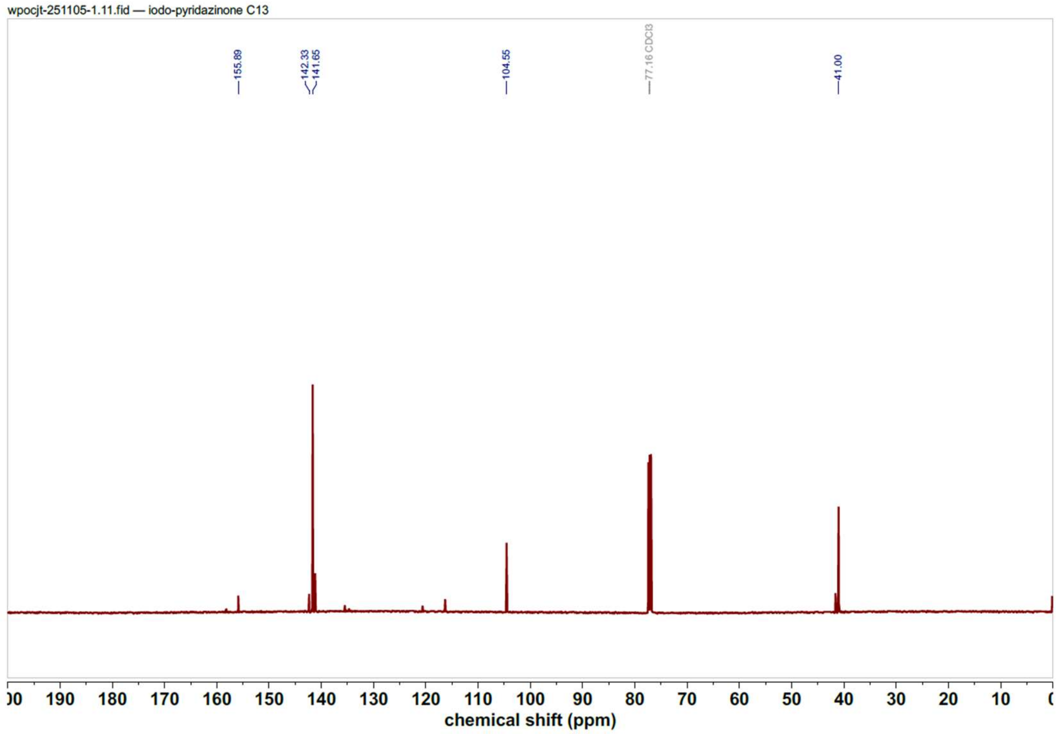


**10**, $^{1}$H NMR (500 MHz, Chloroform-*d*)

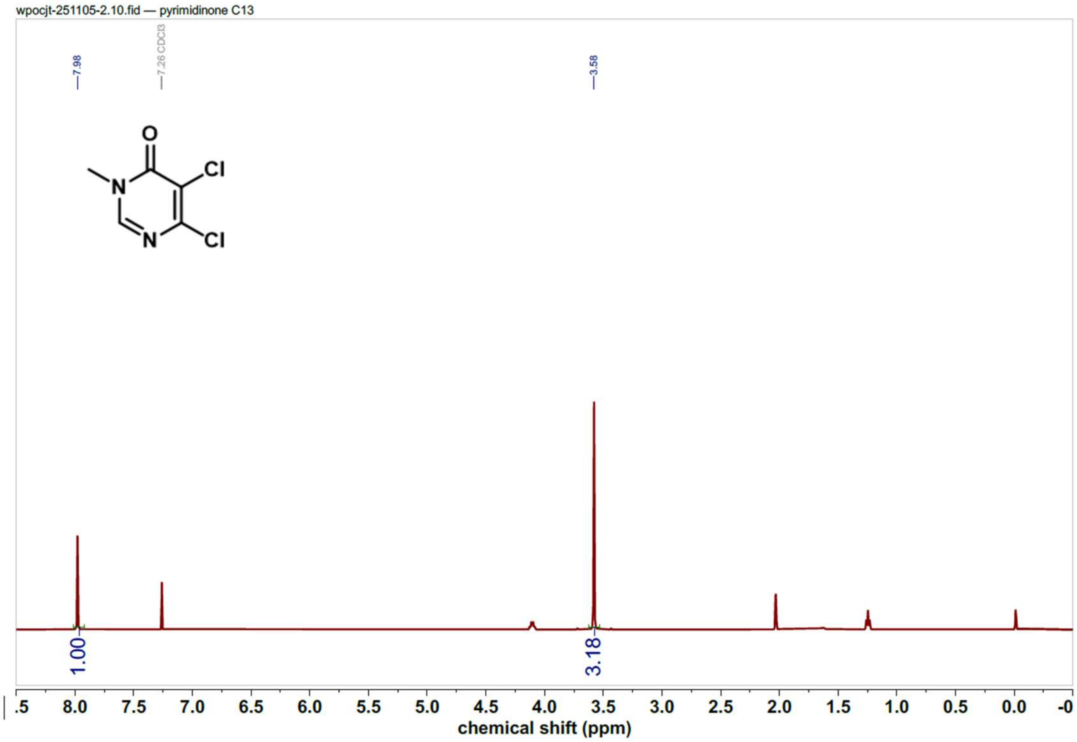


**10**, $^{13}$C NMR (126 MHz, Chloroform-*d*)

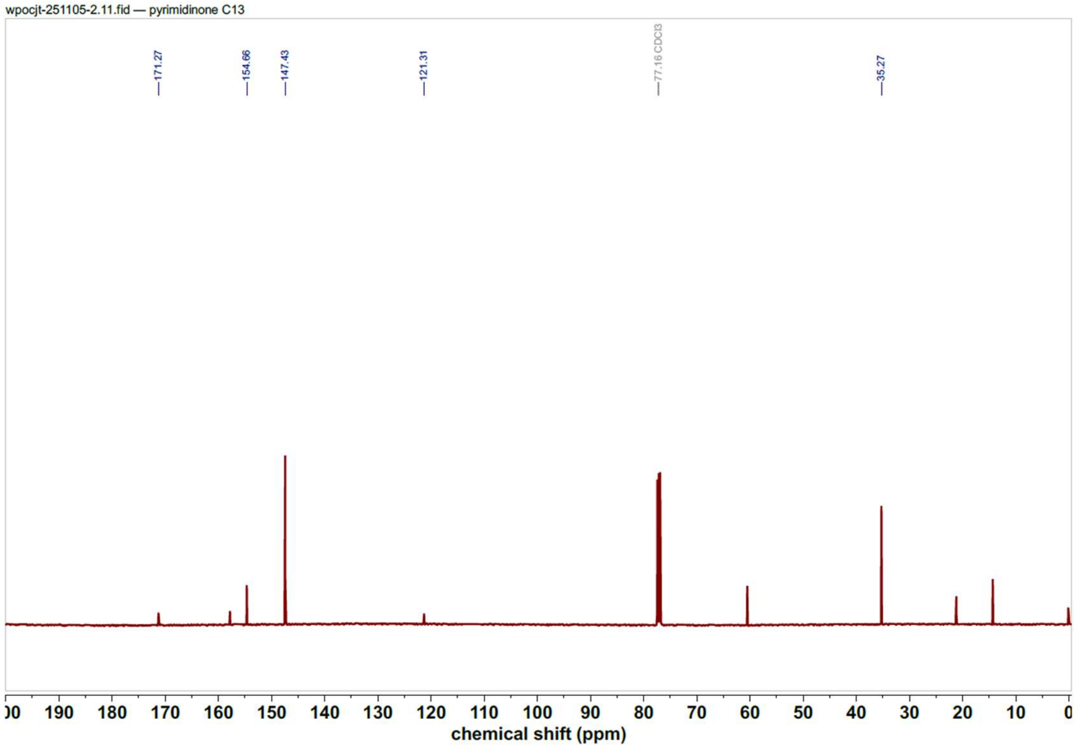


**8**, $^{1}$H NMR (500 MHz, methanol-$d_4$)

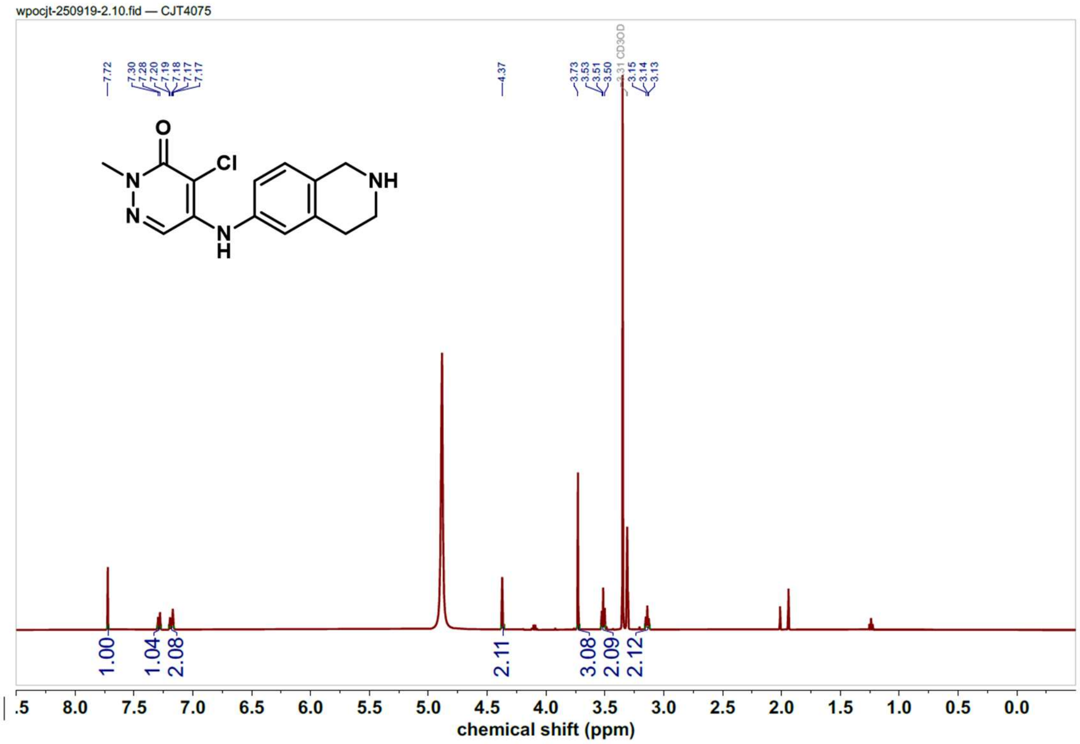


**8**, $^{13}$C NMR (126 MHz, methanol-$d_4$)

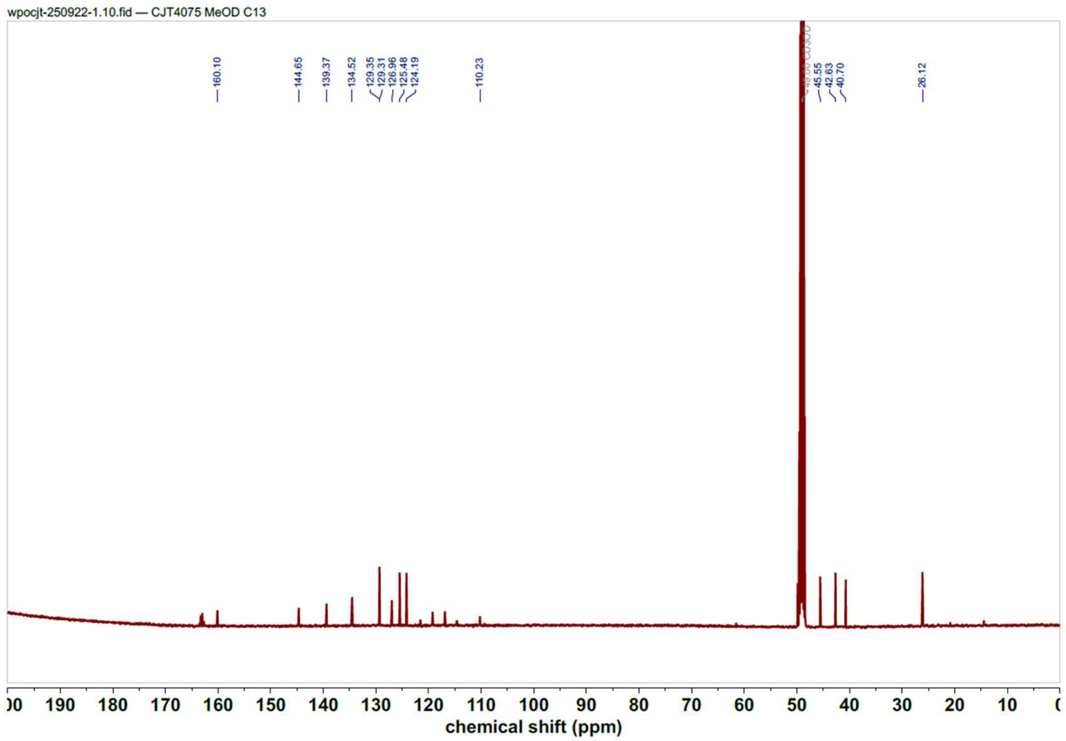


**4**, $^{1}$H NMR (400 MHz, Chloroform-*d*)

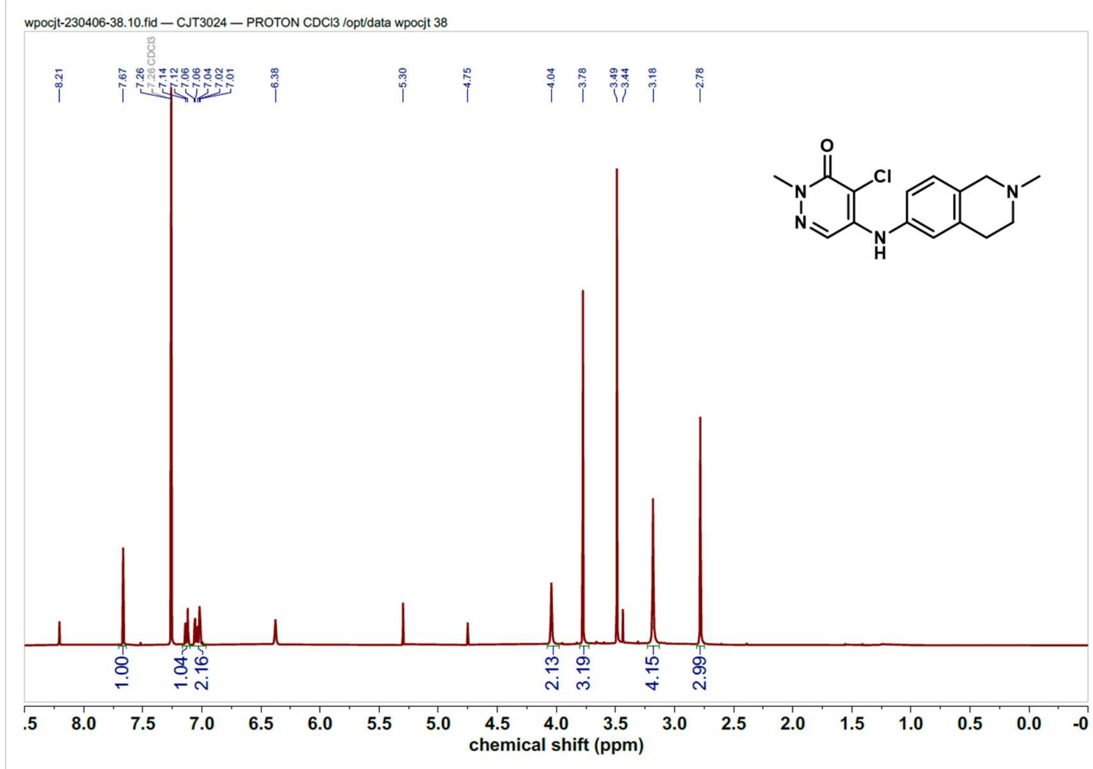


**4**, $^{13}$C NMR (126 MHz, Chloroform-*d*)

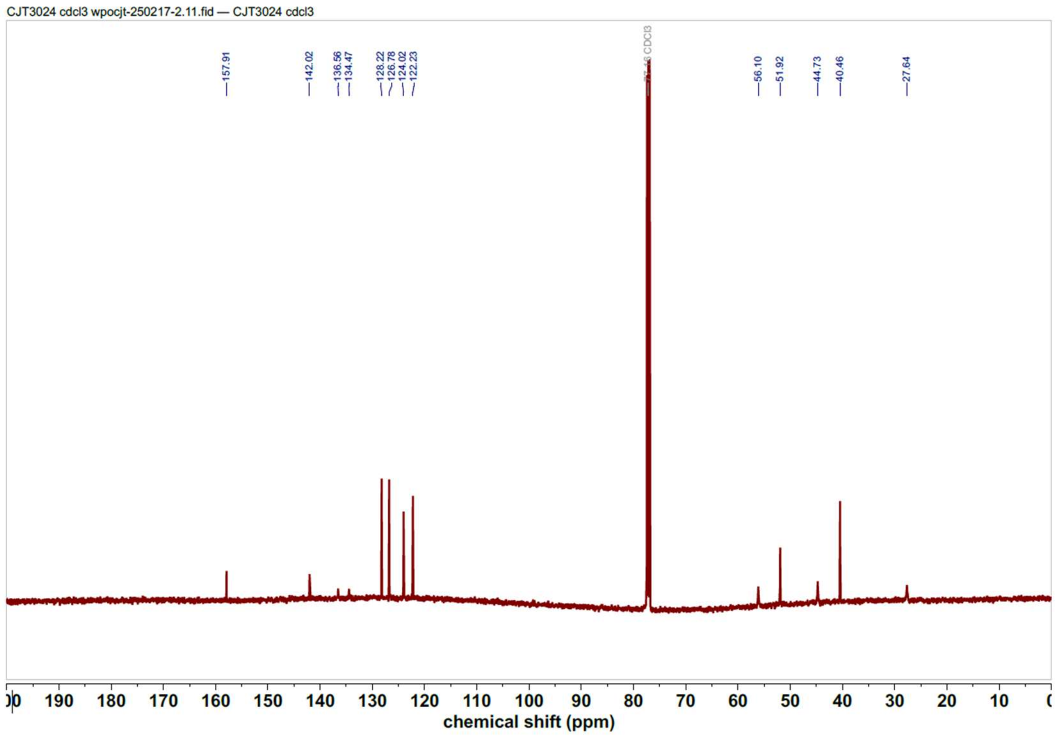


**5**, $^{1}H$ NMR (400 MHz, methanol-$d_4$)

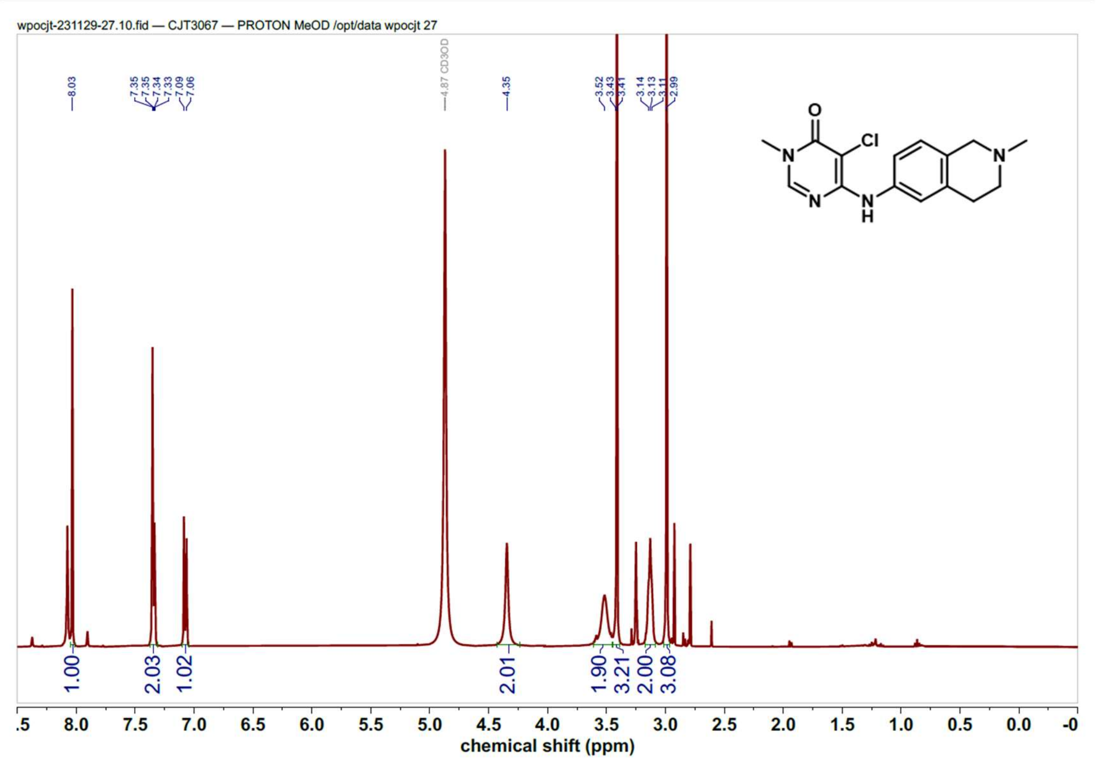


**5**, $^{13}C$ NMR (126 MHz, Chloroform-$d$)

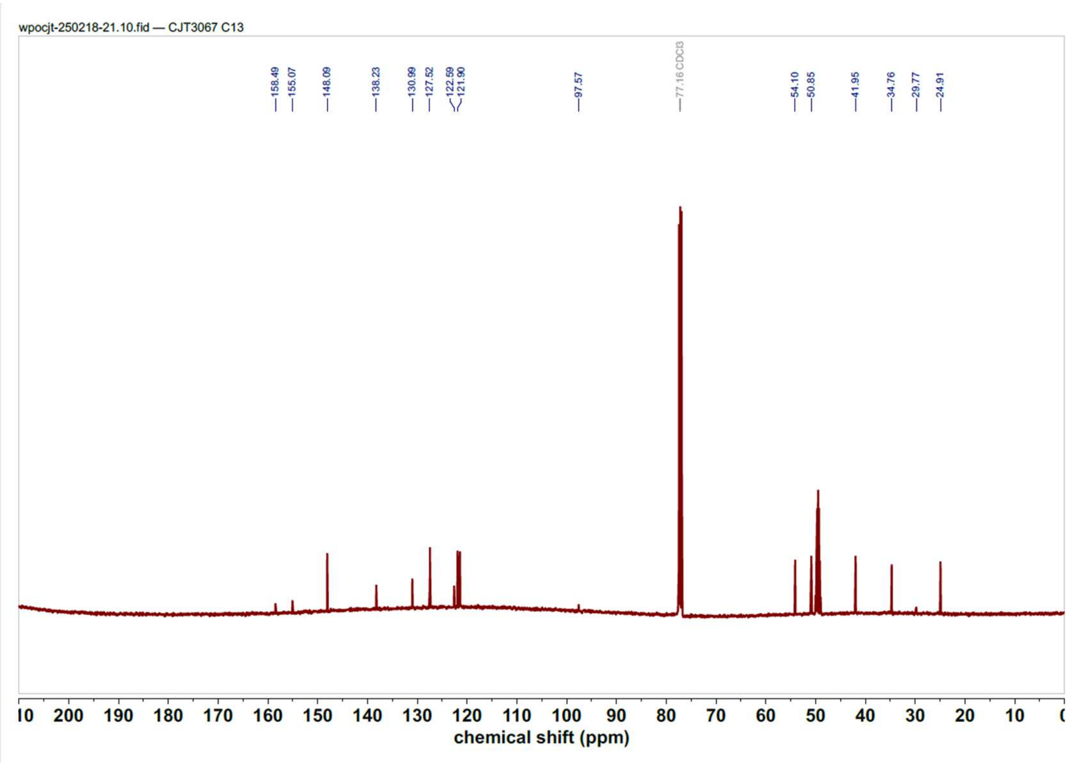


**6**, $^{1}$H NMR (500 MHz, methanol-$d_4$)

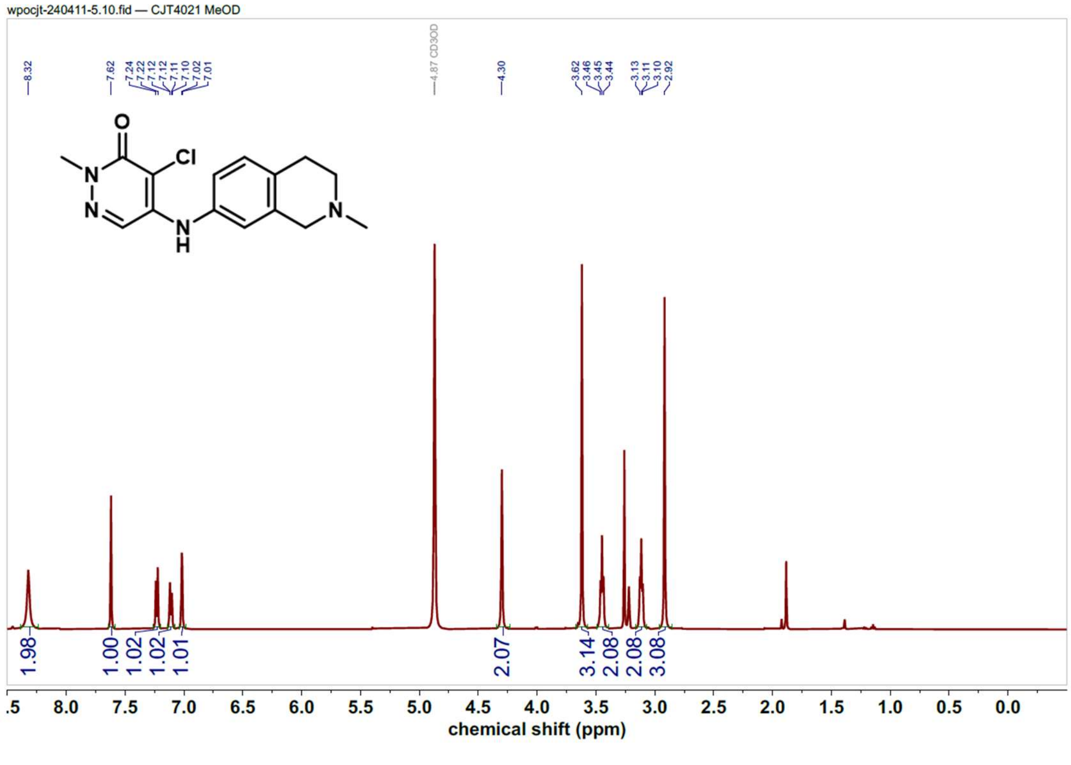


**6**, $^{13}$C NMR (126 MHz, Chloroform-$d$)

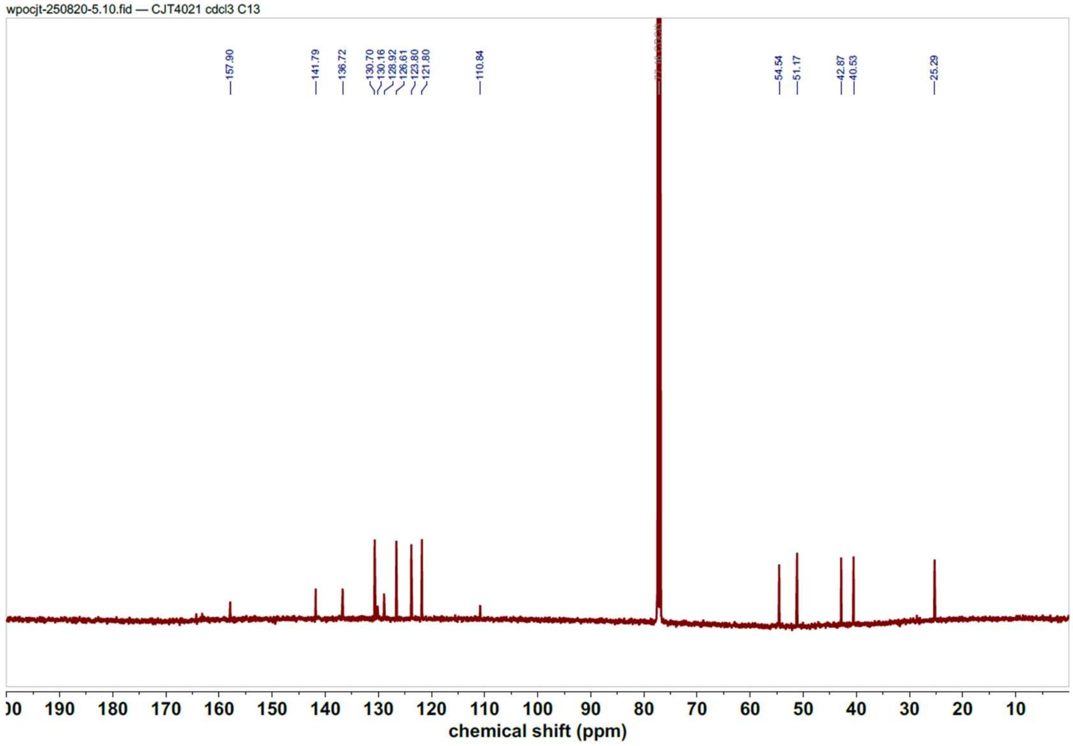


### References


(1) Zahid, H.; Costello, J. P.; Li, Y.; Kimbrough, J. R.; Actis, M.; Rankovic, Z.; Yan, Q.; Pomerantz, W. C. K. Design of Class I/IV Bromodomain-Targeting Degraders for Chromatin Remodeling Complexes. *ACS Chem. Biol.* **2023**, *18* (6), 1278–1293. https://doi.org/10.1021/acschembio.2c00902.

(2) Zahid, H.; Buchholz, C. R.; Singh, M.; Ciccone, M. F.; Chan, A.; Nithianantham, S.; Shi, K.; Aihara, H.; Fischer, M.; Schönbrunn, E.; dos Santos, C. O.; Landry, J. W.; Pomerantz, W. C. K. New Design Rules for Developing Potent Cell-Active Inhibitors of the Nucleosome Remodeling Factor (NURF) via BPTF Bromodomain Inhibition. *J. Med. Chem.* **2021**, *64* (18), 13902–13917. https://doi.org/10.1021/acs.jmedchem.1c01294.
